# Lipidic *α*-synuclein fibrils catalyze the formation of pathological mature fibrils

**DOI:** 10.64898/2026.11.04.764198

**Authors:** Alisdair Stevenson, Jesper E. Dreier, Nina Baumgartner, Natascha T. Christiansen, Céline Galvagnion, Thomas C. T. Michaels

**Author notes:** Authors to whom correspondence should be addressed. Electronic mail.

## Abstract

The aggregation of *α*-synuclein into amyloid fibrils is a hallmark of a collection of neurodegenerative diseases termed synucleinopathies, including Parkinson’s disease (PD). Interactions between *α*-synuclein and lipid membranes modulate *α*-synuclein aggregation and can produce protofibril structures consisting of both protein and lipid, termed lipidic fibrils. These lipidic fibrils can mature into mature fibrils, which resemble *ex vivo* fibrils extracted from Lewy bodies found in post-mortem brain samples of PD patients. Here, through the combination of experiments and kinetic theory, we study the influence of lipids and lipidic fibrils on the formation of mature fibrils. We find that lipidic fibrils catalyze mature fibril formation via a heterogeneous primary nucleation mechanism, which lowers the activation energy barrier of mature fibril formation relative to lipid-independent primary nucleation. The final stage of the reaction is governed by mature fibril elongation becoming limited by the rate at which protein monomers dissociate from the metastable lipidic fibrils, confirming that fibrils form in order of stability following Ostwald’s Rule of Stages. This mechanism is validated using the small molecule inhibitor ambroxol, which selectively suppresses the formation of lipidic fibrils and in this manner shifts the aggregation kinetics to the slow, lipid-independent pathway. Taken together, these findings reveal that lipidic fibrils act as on-pathway intermediates and accelerate the aggregation of mature fibrils, which could provide a quantitative framework to design novel therapeutic strategies to inhibit pathological *α*-synuclein aggregation.

**SIGNIFICANCE STATEMENT:** Amyloid aggregation is associated with a broad range of diseases such as Parkinson’s disease, where Lewy Bodies are predominantly formed of *α*-synuclein fibrils but are also rich in lipids. The influence of lipids on the formation of mature fibrils is currently unknown. In this work, we present a global kinetic model that, when integrated with experimental data, reveals the mechanistic landscape of lipid-dependent *α*-synuclein mature fibril formation. Crucially, the metastable lipidic fibrils that initially form catalyze the nucleation of mature fibrils that resemble *ex vivo* fibrils from post-mortem brain samples. These results provide a quantitative framework for understanding how lipids alter the energetic landscape of *α*-synuclein aggregation and offer a strategy for the targeted disruption of lipid-dependent amyloid aggregation.

## I. INTRODUCTION

Ostwald’s Rule of Stages states that a system undergoing a phase transition toward a state of higher stability does not necessarily reach the global thermodynamic minimum directly, but instead progresses through a sequence of intermediate states with the smallest loss of free energy that are, therefore, kinetically more favorable to form^1^. The phenomenon of navigating through transient phases to reach the minimum energy state is observed in systems as diverse as calcium carbonate precipitating from water, the self-assembly of Boc-FF peptides and, of particular relevance to this study, the aggregation of *α*-synuclein to form amyloid fibrils in the presence of lipids^2–4^. Amyloid fibrils are associated with a diverse range of diseases^5–8^, where a hallmark of Parkinson’s disease (PD) and related synucleinopathies is abnormal proteinaceous deposits primarily composed of *α*-synuclein fibrils^9^. *α*-synuclein is an intrinsically disordered protein with a native function associated with synaptic vesicle trafficking and lipid membrane remodeling, both of which depend on proteinlipid interactions^10–16^. Furthermore, the proteinaceous deposits found in PD have also been found to contain high concentrations of lipids^6,9–12^. The transition of *α*-synuclein from a functional monomer to the pathologically associated amyloid fibril state is crucial to PD pathogenesis and, therefore, elucidating the microscopic reaction steps governing the aggregation of *α*-synuclein has motivated significant research efforts in recent decades^17–20^.

*α*-synuclein is relatively resistant to aggregation under quiescent conditions and neutral pH *in vitro*^18,19^, likely due to slow homogeneous fibril nucleation. One way to overcome this slow nucleation rate is to aggregate *α*-synuclein fibrils in the presence of lipids^4,14,15,21–27^. Although *α*-synuclein can bind to various biologically relevant lipids, such as cardiolipin and gangliosides, not all lipid membranes induce aggregation *in vitro*^26,28,29^. For the anionic lipid 1,2-dimyristoylsn-glycero-3-phospho-L-serine (DMPS), we previously determined the mechanistic pathway of lipid-induced *α*-synuclein aggregation through the use of chemical kinetics theory^27,30^.

Inducing *α*-synuclein aggregation with DMPS produces fibrils with structural characteristics distinct from mature fibrils formed in the absence of lipids^4,18,24,31,32^. Critically, these fibrils are formed via the coaggregation of lipids and protein monomers to produce lipidic fibrils^4,24,26,31–34^. The direct incorporation of lipids into the fibril structure, as confirmed with Small-Angle X-Ray Scattering and Neutron Scattering (SAXS and SANS, respectively) and Nuclear Magnetic Resonance (NMR)^26,35^, results in lipidic fibrils that are more flexible and shorter in negative stain electron microscopy (EM) images compared to mature fibrils formed only of *α*-synuclein monomers^27,36^. Furthermore, *α*-synuclein undergoes induced folding upon binding to the lipid surface, where the formation of an *α*-helix structure in the N-terminal domain exposes the NAC domain, whose hydrophobic residues drive aggregation^24,37^. We have previously demonstrated that lipidic fibrils are a metastable intermediate phase in the rugged energy landscape of *α*-synuclein structural forms^4^. When lipidic fibrils are heated, the incorporated lipids are shed to produce the more thermodynamically stable mature fibril structure (consisting only of protein) that resembles *ex vivo* fibrils extracted from Lewy bodies found in post-mortem brain samples of PD patients^38^.

These observations raise the question of how the presence of lipids and formation of lipidic fibrils interplay with the formation of mature fibrils. Here, we combine aggregation kinetics experiments with chemical kinetics theory to rationalize the accelerated lipid-dependent aggregation pathway forming mature *α*-synuclein fibrils that follows a double-sigmoid kinetic profile. We find that the metastable lipidic fibrils are onpathway intermediates that accelerate the aggregation of mature fibrils through a heterogeneous primary nucleation reaction. This heterogeneous nucleation pathway is the kinetically more accessible mechanism of forming mature fibrils, as it reduces the activation energy barrier by *∼* 25 kJmol^*−*1^ relative to lipid-independent primary nucleation in our experimental setup. As the lipidic fibrils undergo dissolution once the thermodynamically more stable mature fibrils are present, our analysis confirms that the different fibril states form in order of increasing stability and follow Ostwald’s Rule of Stages. Through the selective inhibition of the formation of lipidic fibrils with the small molecule ambroxol, we confirm that the acceleration observed in the lipid-dependent pathway of mature fibril formation can be attenuated, suggesting a promising novel therapeutic strategy to control aberrant *α*-synuclein aggregation facilitated by lipids.

## II. RESULTS

### Lipid-dependent aggregation pathway follows a double sigmoid kinetic profile

In PEGylated plates in the absence of lipids, the spontaneous nucleation of mature *α*-synuclein fibrils is too slow to observe on experimentally accessible timescales. Therefore, we utilized polystyrene (PS) microwell plates, as PS induces *α*-synuclein aggregation under conditions where aggregation is not observed otherwise, to study the effects of lipids on mature fibril aggregation. PS surfaces have been found to induce the aggregation of *α*-synuclein to form mature fibrils (red curve Fig. 1a), where it is proposed that PS induces primary nucleation^39–43^. *α*-synuclein has a high affinity for PS surfaces, binding primarily at the N-terminal domain, with a *K*_*d*_ *≈* 2 *nM*^44^, where the N-terminal and central domains of *α*-synuclein adopt an *α*-helical conformation upon binding to the PS surface^45,46^. The increased propensity to aggregate in the presence of PS is likely due to the induced folding upon binding to the PS surface and stabilization of fibril nuclei that form at the PS surface by reducing surface exposure to the bulk solvent. The effect of PS on *α*-synuclein aggregation is not only relevant to *in vitro* studies. PS is an abundant plastic with many applications and, as a result, is a major component of environmental micro- and nanoplastics^47^. Furthermore, micro- and nanoscale PS can cross the blood-brain barrier and has been identified in post-mortem brain tissue^48^, where the implications of this on accelerating *α*-synuclein aggregation *in vivo* are currently unknown. The product of PS-induced aggregation is pure *α*-synuclein mature fibrils characterized by a straight, rigid morphology (Fig. 1b).

**FIG. 1.**
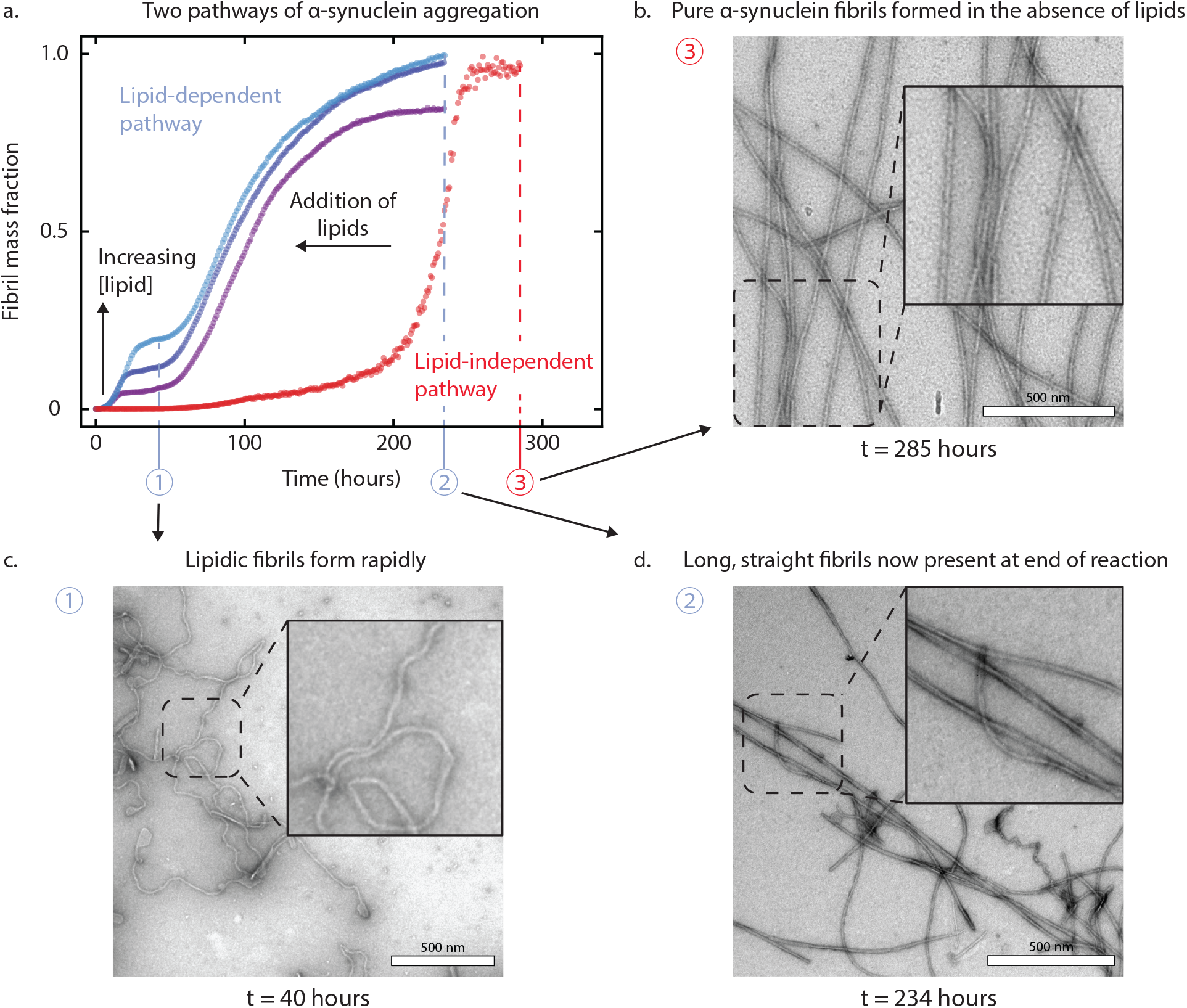
Lipid-dependent *α*-synuclein aggregation in PS plate displays a double sigmoid kinetic profile with distinct fibril product morphology. **(a)** Experimental ThT fluorescence measurements of the *in vitro* kinetics of *α*-synuclein aggregation in the absence or presence of lipids in the PS plate assay. Lipid-independent *α*-synuclein aggregation exhibits a single sigmoid kinetic profile, with a relatively long lag phase, where the nucleation of mature fibrils is induced by the presence of the PS surface (red curve). Upon the addition of DMPS lipids, the aggregation reaction follows an accelerated double sigmoid kinetic profile. An initial protein concentration of *m*_tot_ = 50 *µ*M of *α*-synuclein and lipid concentration of *L*_tot_ = 100 (purple), 200 (blue) and 400 *µ*M (light blue). **(b)** Negative-stain EM images of fibrils produced by the lipid-independent *α*-synuclein aggregation pathway show relatively long fibrils with a straight, rigid morphology formed only of protein monomers. An initial protein concentration of 50 *µ*M of *α*-synuclein was used to form the mature fibrils. **(c)** EM images of fibrils isolated after the first growth phase (*t* = 40 hours) in the lipid-dependent aggregation pathway show lipidic fibrils, with a more flexible morphology, formed by the coaggregation of protein monomers and lipids. An initial protein concentration of 50 *µ*M of *α*-synuclein and a lipid concentration of 400 *µ*M of DMPS is used to form the lipidic fibrils. **(d)** EM images of fibrils after the second growth phase (*t* = 234 hours) in the lipid-dependent aggregation pathway show morphological features consistent with mature fibrils.

Using the PS plate assay, we sought to elucidate the effect of the anionic phospholipid DMPS on *α*-synuclein aggregation. DMPS is known to modulate *α*-synuclein aggregation in PEGylated plates, acting as either an accelerator or an inhibitor depending on the lipid-to-protein ratio (*r* = *L*_tot_*/m*_tot_)^24,27^. In the presence of DMPS lipids, we observed a double-sigmoid Thioflavin T (ThT) fluorescence profile, consisting of two well-defined growth phases separated by an intermediate plateau (Fig. 1a). To elucidate the nature of this complex double-sigmoid kinetic profile, referred to here as the lipid-dependent aggregation pathway, we first validated that the initial growth phase corresponds to the rapid coaggregation of lipidic fibrils. EM confirmed a clear morphological difference in the fibrils present at the end of the first and second growth phases (Fig. 1c and 1d, respectively). Images obtained at the intermediate plateau (the end of the first growth phase) reveal short, flexible fibrils that are morphologically similar to the lipidic fibrils formed in PEGylated plates^27^. In contrast, after the second growth phase, we find a homogeneous population of relatively long, straight structures, resembling the morphology of mature fibrils formed in the absence of lipids (Fig. 1d and 1b, respectively). It is possible that the fibril product from the second growth phase is still composed of both protein monomers and lipids, potentially with a lower stoichiometry of lipids per protein monomer to produce the more rigid fibrils observed. However, we assume the second growth phase corresponds to the aggregation of mature fibrils formed only of protein monomers and is not dependent on lipids, given the morphological resemblance in fibrils shown in Fig. 1d and 1b.

### First growth phase corresponds to lipidic fibril formation

The first growth phase retains the key qualitative hall-marks of lipidic fibril aggregation observed in PEGylated plates, including the characteristic switch from a lipid-to a protein-dependent lipidic fibril mass yield when either varying the initial lipid (Fig. 2a) or protein monomer concentration (Fig. 2b)^27^. To understand whether the presence of the PS surface modulates the established mechanisms of lipidinduced aggregation and its kinetics, we applied a previously developed chemical kinetics model of lipid-induced protein aggregation to the first growth phase of the kinetic profiles (Fig. 2c and Fig. 2d)^27,30^. The details of the kinetic model describing lipidic fibril aggregation can be found in the Methods section (Eq. (1)). In short, the model describes the time evolution of the aggregation reaction in terms of the fractional coverage of the lipid surface by protein monomers and lipidic fibrils, rather than their bulk concentrations that is typically used in kinetic descriptions of aggregation in the absence of lipids^49–55^. Protein binding to the lipid surface is treated as a rapid pre-equilibrium, set by the dissociation constant *K*_*D*_ and the number of lipids bound per monomer, *β* . Lipidic fibrils form through two-step primary nucleation: oligomers form from bound and free monomers (rate constant *k*_*o*_, reaction orders *n*_1_ and *n*_2_) and then convert into elongation-competent fibrils (rate constant *k*_*c*_). These fibrils grow by adding lipidbound monomers (rate constant 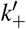), a process that consumes lipids at a stoichiometry of *α* lipids per protein monomer incorporated into the lipidic fibril. The model yields analytical integrated rate laws, Eq. (1), which predict that the final fibril mass is limited by lipid for lipid-to-protein ratios *r* = *L*_tot_*/m*_tot_ *< α* and by protein for *r ≥ α*.

**FIG. 2.**
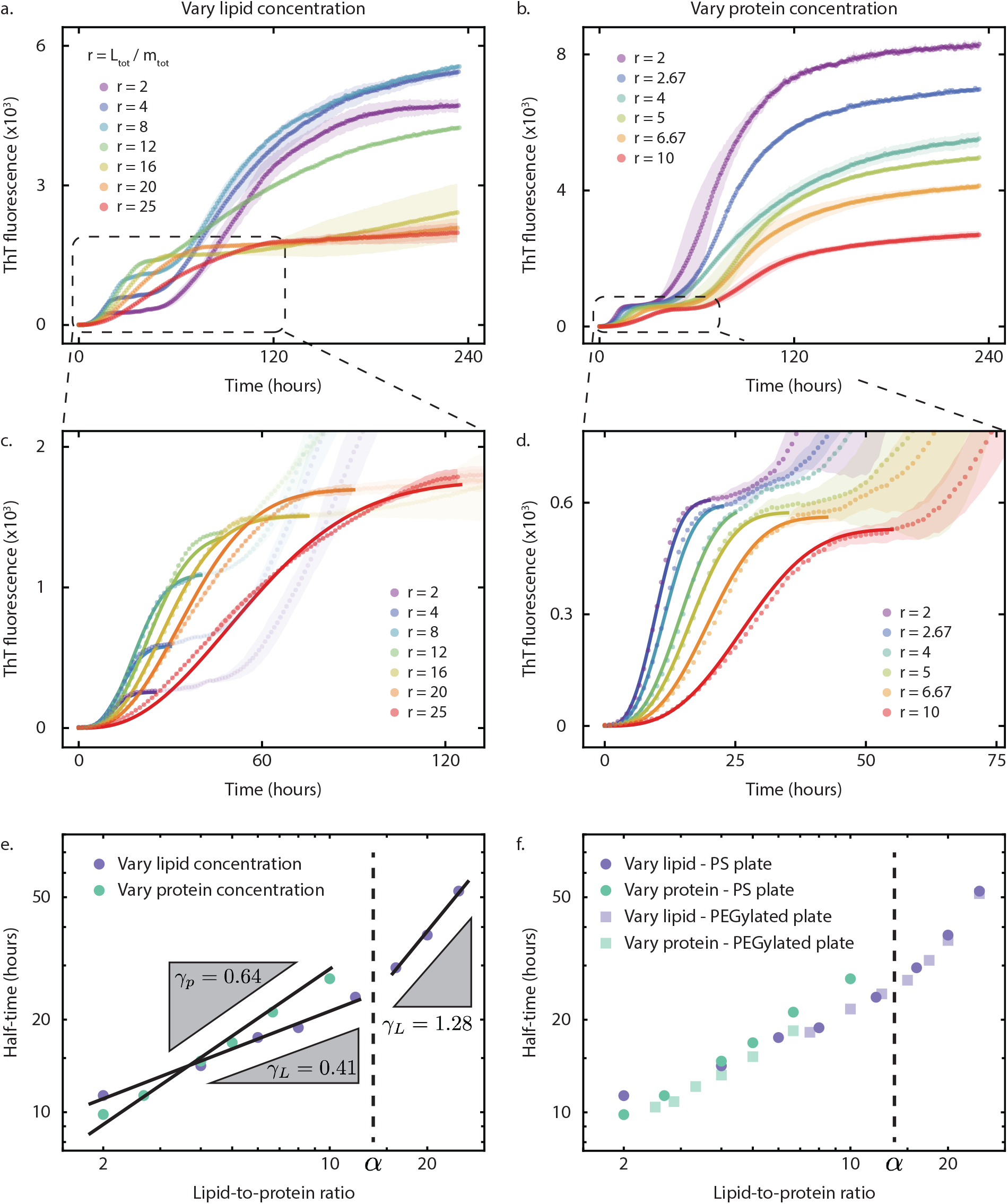
First growth phase of lipid-dependent *α*-synuclein aggregation corresponds to the formation of lipidic fibrils. Experimental aggregation curves monitoring the kinetics of fibril formation using ThT fluorescence with **(a)** constant *m*_tot_ = 50 *µ*M and varying *L*_tot_ = 100, 200, 400, 600, 800, 1000 and 1250 *µ*M and **(b)** varying *m*_tot_ = 20, 30, 40, 50, 75 and 100 *µ*M and constant *L*_tot_ = 200 *µ*M. **(a)** For *r < α* aggregation curves, where *α* = 14 for all fits, the yield of lipidic fibrils is lipid-limited and, therefore, protein monomers are available for the second growth phase. The curves are mean values of triplicates, with the shaded regions representing the standard deviation. For *r > α* aggregation curves, the yield of lipidic fibrils is now protein-limited and, therefore, the protein monomer concentration is sufficiently low to not observe a consistent second growth phase. **(b)** As *r < α* for all curves, the yield of lipidic fibrils is lipid-limited and, therefore, protein monomer is available for the second growth phase. Isolating the first sigmoid and globally fitting our integrated rate law, Eq. (1), describing <u>lipidi</u>c fibril aggregation produces quantitatively accurate fits when varying **(c)** lipid and **(d)** protein monomer concentrations (average 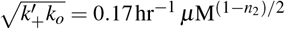 and 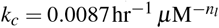, *n*_*l*_ = 0.21, and constant *n*_1_ = 0 and *n*_2_ = 1.5). Reaction orders *n*_1_ and *n*_2_ were taken from the values determined from *t*_1*/*2_ scaling analysis (Fig. 2e). **(e)** The half-time of aggregation reveals the reaction orders through scaling analysis of our theoretical model, where we identify *n*_1_ = 0 and *n*_2_ = 1.5 for the dependence of oligomer formation on bound and free protein monomers, respectively. **(f)** Comparison of half-time data for kinetic profiles measured in polystyrene and PEGylated plates confirms consistency in macroscopic kinetic behavior of lipidic fibril aggregation^27^.

To constrain the global fitting of the kinetic model to the experimental data, we fixed as many parameters as possible independently of the kinetic fits. The number of lipids bound per protein monomer when the membrane is fully covered, *β* = 28.2*±* 0.8, was taken from previous experimental measurements^24^. The lipid-to-protein stoichiometry in lipidic fibrils, *α* = 14, was determined by comparing the plateau ThT values of the first growth phase as a function of *r* with the predicted steady-state behavior, Eq. (13). The reaction orders, *n*_1_ and *n*_2_, were identified from the scaling of the half-time (*t*_1*/*2_) of the first growth phase with lipid and protein concentration (Fig. 2e). We previously revealed that *t*_1*/*2_ follows a scaling relation depending on whether *L*_tot_ or *m*_tot_ is being varied^27,30^. Analysis of *t*_1*/*2_ in the first growth phase revealed the reaction orders associated with the oligomer nucleation process as *n*_1_ = 0 and *n*_2_ = 1.5 (Fig. 2e), identical to those in PEGylated plates^27^. This leaves only 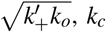 and *n*_*l*_, where the lipid dependence of oligomer conversion is described by *n*_*l*_. We determined these parameters by globally fitting the integrated rate law to the full set of kinetic profiles, yielding 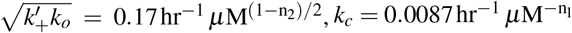,and *n*_*l*_ = 0.21. The model accurately describes the first growth phase across all lipid and protein concentrations (Fig. 2c and 2d). Notably, the half-times in the PS plate are indistinguishable within experimental uncertainty from those measured in PEGylated plates (Fig. 2f)^27^. These findings demonstrate that the first growth phase in the double sigmoid kinetic profiles corresponds to the lipid-induced aggregation of *α*-synuclein, producing lipidic fibrils, and that the PS surface does not perturb the mechanistic pathway or macroscopic kinetics of lipidic fibril aggregation.

### Lipidic fibrils catalyze the formation of mature fibrils

The observed aggregation kinetics suggest the lipidic fibrils are accelerating the formation of mature fibrils, as the introduction of lipids decreased the overall timescale of aggregation when compared to the lipid-independent kinetic profiles (Fig. 1a). This acceleration suggests that the lipids or lipidic fibrils, or both, are facilitating a more rapid transition from the monomeric to mature fibril form. Alternatively, the metastable lipidic fibrils could act as a kinetic trap for *α*-synuclein monomers, where the formation of either lipidic or mature fibrils represent distinct pathways of aggregation in the rugged energy landscape of *α*-synuclein structural states. In this scenario, the distinct fibril species would compete for the shared pool of protein monomer and the protein monomers consumed to form lipidic fibrils would then be no longer available to the lipid-independent pathway, and the dynamics driving the second growth phase would be slowed. The reduced protein pool would, therefore, produce a double sigmoid profile with a slower overall timescale of aggregation with respect to the aggregation of mature fibrils in the absence of lipids, which was not observed in the data.

To elucidate the mechanism underlying the accelerated formation of mature fibrils in the presence of lipids, we analyzed the duration of the intermediate plateau, *t*_lag_, as a function of *r* (Fig. 3a). At low *r* values, where lipidic fibril yield is limited by lipid availability, a fraction of the initial protein monomer pool remains in solution after the completion of the first growth phase. We found that *t*_lag_ is inversely correlated with *r* and, therefore, the initial lipid concentration in the lipid-limited regime of lipidic fibril yield. As the initial lipid concentration is increased, the yield of lipidic fibrils increases and the pool of free protein monomers remaining after the first growth phase is reduced, causing *t*_lag_ to decrease. The inverse correlation between the concentration of protein monomer remaining after lipidic fibrils have formed and *t*_lag_ suggests that the second growth phase is initiated by a nucleation process dependent on the presence of lipidic fibrils, rather than solely on the protein monomers. Indeed, a kinetic model including a heterogeneous nucleation rate proportional to the lipidic fibril mass concentration (at the end of the first growth phase) predicts that the lag time of the second growth phase scales as 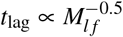, in line with the experimental data (Fig. 3a, bottom panel).

**FIG. 3.**
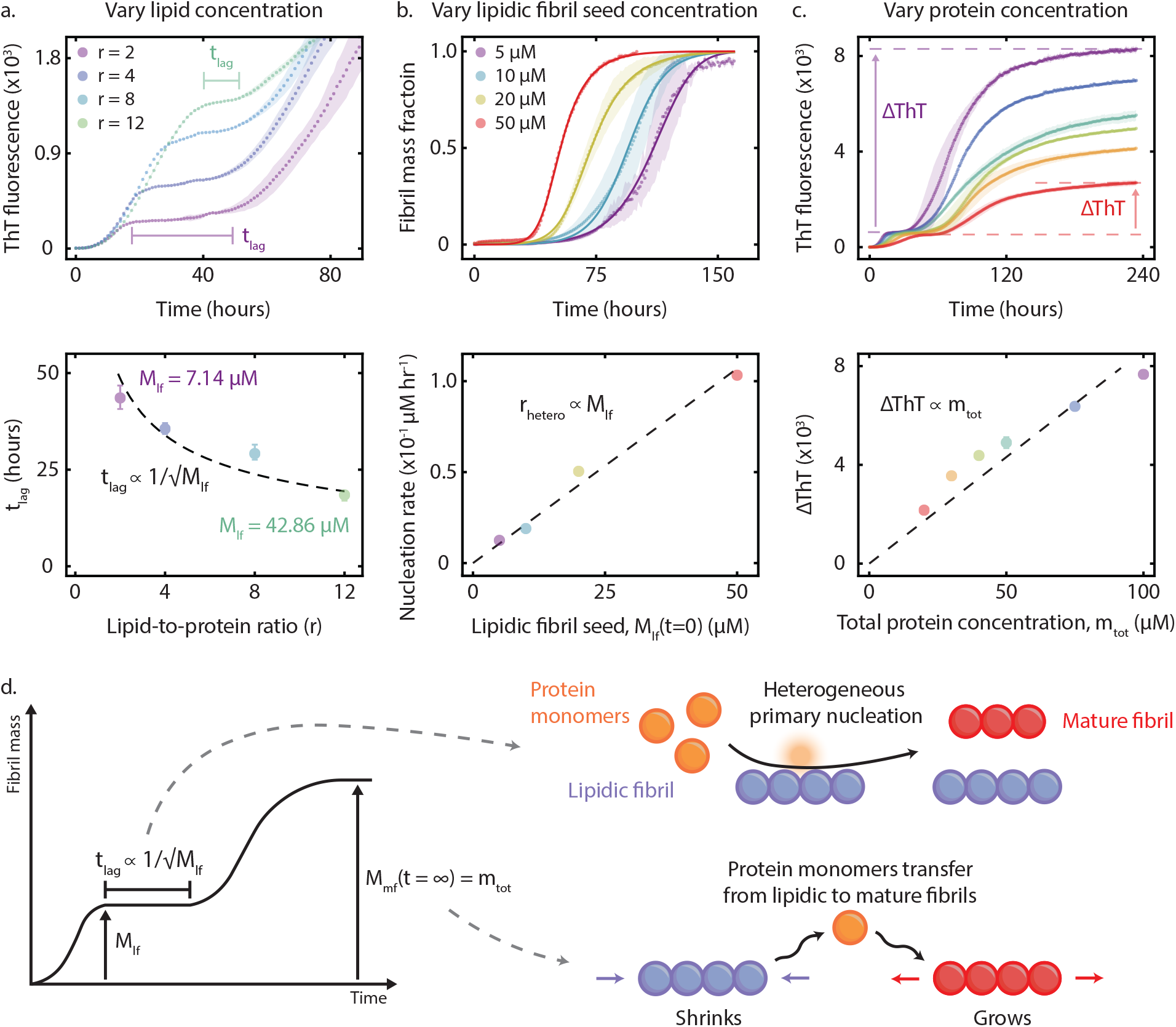
Macroscopic features of kinetic profiles reveal microscopic insight into the mechanistic pathway of lipid-dependent *α*-synuclein aggregation. **(a)** The length of the intermediate plateau, *t*_lag_, decreases with increasing lipidic fibril mass concentration, *M*_*l f*_ . As more protein monomer is consumed in the first growth phase to produce lipidic fibrils, the nucleation event initiating the second growth phase must depend on the concentration of lipidic fibrils present. Experimental conditions are described in Fig. 2. **(b)** Top panel: The presence of increasing concentrations of lipidic fibril seeds in the PS plate accelerates mature fibril aggregation but the kinetic profiles still display prolonged lag phases, confirming a nucleation event forming mature fibrils is an essential feature of the mechanistic landscape. Bottom panel: Individually fitting a coarse-grained primary nucleation rate in our theoretical model (solid lines in top panel), Eq. (6), to capture the time evolution of mature fibril aggregation reveals a direct linear proportionality of the primary nucleation rate on *M*_*l f*_ . **(c)** The direct linear proportionality of ΔThT on *m*_tot_ confirms that all protein monomers are consumed to form mature fibrils (see Supplementary Material Fig. S1 for similar analysis of vary lipid dataset). **(d)** Schematic representations of the novel microscopic reaction steps that feature in the mechanistic pathway of lipid-dependent *α*-synuclein aggregation. Heterogeneous primary nucleation of mature fibrils involves protein monomers forming mature fibril nuclei that are stabilized at the surface of lipidic fibrils. The dissolution of lipidic fibrils, once mature fibrils are present, eventually converts all protein mass in the system to mature fibril form.

The inclusion of heterogeneous primary nucleation involving lipidic fibrils is further supported by the kinetic profiles at higher *r* values (Fig. 2a). As *r* increases, a larger proportion of the protein monomer pool is consumed to form lipidic fibrils during the first growth phase. Crucially, when *r* is sufficiently high to effectively sequester the entire protein monomer pool in lipidic fibril form, the second growth phase is suppressed. The absence of a second growth phase in this case, despite a high concentration of lipidic fibrils, confirms that the lipidic fibrils act as a scaffold requiring a critical concentration of free protein monomer for the nucleation of mature fibrils. Taken together, the data confirm a nucleation barrier exists between these two fibril species rather than the transition between aggregated forms being a direct unimolecular structural rearrangement of lipidic to mature fibrils, that could occur without free protein monomers present, and would not produce a distinct intermediate plateau between growth phases.

We further investigated the nucleation-controlled intermediate plateau with a series of seeding experiments using preformed lipidic fibrils (Fig. 3b). The lipidic fibril seeds were produced in the PS plate, with *m*_tot_ = 100 *µ*M and *L*_tot_ = 800 *µ*M, where samples were taken from this solution at the end of the first growth phase (tracked with ThT fluorescence). We varied the concentration of lipidic fibril seeds and observed that *t*_1*/*2_ decreased from *∼*110 hours to *∼*45 hours upon a 10-fold increase in seed concentration (Fig. 3b top panel). We fitted these kinetic curves to a secondary nucleation-dominated kinetic model, with an individually fitted rate of heterogeneous primary nucleation. The extracted rate of primary nucleation was found to be linearly dependent on lipidic fibril mass concentration (Fig. 3b bottom panel), in line with the conclusion from Fig. 3a. Alternatively, with a fixed seed concentration, a 10-fold increase in the free protein monomer concentration led to a reduction in *t*_1*/*2_ from *∼*125 hours to *∼*60 hours. Despite the reduction in *t*_1*/*2_, the fitting analysis revealed that the nucleation rate did not depend on protein monomer concentration, suggesting a saturated contribution of protein monomers to the heterogeneous primary nucleation reaction flux (Supplementary Material Fig. S2). Critically, in all seeding conditions, a distinct lag phase persisted before the onset of the sigmoidal kinetic profiles tracking the aggregation of mature fibrils. The inability of lipidic fibril seeds to eliminate the lag phase, contrasting with the rapid increase in ThT fluorescence observed when mature fibril seeds are used (Supplementary Material Fig. S3)^36^, confirms that a new nucleation event is required before the aggregation reaction can proceed (mature fibril propagation driven by lipid-independent elongation). The different seeding capacity of lipidic and mature fibrils provides further evidence, along with EM characterization, that these are structurally distinct fibril species. Taken together, these results support our description of a heterogeneous primary nucleation reaction, positioning the lipidic fibrils as a catalytic surface that facilitates the formation of the structurally distinct mature fibrils, given the different seeding capacity, from the remaining protein monomer pool. As *r < α* when producing the lipidic fibril seeds, the yield of lipidic fibrils is lipid-limited, leaving excess protein monomer in the solution that is transferred to the seeding experiments. To avoid affecting the seeding capacity of the lipidic fibrils, the solution was not washed or spun down to try to remove the excess protein monomer. Therefore, we account for the excess free protein monomers that are also transferred along with lipidic fibril seeds. Our analysis of the data presented in Fig 3a and 3b quantitatively demonstrates that the catalytically active species in the mechanistic landscape of lipid-dependent *α*-synuclein aggregation is the lipidic fibrils and not the DMPS lipids alone. A physically realistic alternative is that the lipid surface provides an energetically favorable pathway for the nucleation of lipidic fibrils as well as mature fibrils. However, as the yield of lipidic fibrils increases in Fig. 3a, *t*_lag_ decreases, despite all of the lipids being incorporated into the lipidic fibril structure, and the increasing concentration of lipidic fibrils corresponds to an increased heterogeneous primary nucleation of mature fibrils in the seeding experiments (Fig. 3b), providing quantitative evidence contradicting this alternative interpretation.

We propose that this mechanism is analogous to surface-catalyzed secondary nucleation, where the lipidic fibrils provide a binding surface that lowers the energy barrier for the formation of the more thermodynamically stable mature fibrils, and term this mechanism heterogeneous primary nucleation. This differs fundamentally from secondary nucleation, as the lipidic fibril surface acts as a catalyst facilitating the formation of a distinct fibril species rather than an autocatalytic propagation of the same species. The acceleration is due to heterogeneous nucleation pathways being more energetically favorable, given the reduced surface exposure of the nucleus that is bound to the heterogeneous surface, compared to homogeneous pathways where nuclei form in solution with their surface fully exposed to solution. These surfaces, such as metal ions and lipids, have been found to induce the aggregation of proteins such as islet amyloid polypeptide and amyloid-*β* 42 (A*β* 42)^56–58^. The acceleration of aggregation in the presence of amyloid fibril intermediates has also been observed *in vitro* for mixed systems of *α*-synuclein and A*β* 42, where A*β* 42 oligomers were specifically identified as the driver of significantly accelerated *α*-synuclein aggregation via a heterogeneous primary nucleation process^59–61^.

### Lipidic fibrils are consumed during the aggregation of mature fibrils

We next investigated whether at the end of the reaction the lipidic fibrils coexist with mature fibrils in a metastable steady state or if the protein pool is entirely converted into the more thermodynamically stable mature fibril form. Morphological characterization via EM supports the latter scenario, as the fibrils imaged at the first plateau share the indicative features of lipidic fibrils (Fig. 1c)^27^, whereas images at the end of the reaction (second plateau) reveal a homogeneous population of straight fibrils, similar to the mature fibrils formed in the absence of lipids (Fig. 1d and 1b, respectively). If lipidic fibrils remain as a metastable state in the system, the protein monomer pool would be reduced for the second aggregation reaction that forms mature fibrils, and we would expect that the increase in ThT fluorescence during the second growth phase (ΔThT) to have a direct linear dependence on the protein monomer concentration remaining. Although ΔThT correlates linearly with the residual monomer concentration, there is a significant non-zero intercept (Supplementary Material Fig. S4). Instead, we find ΔThT has a direct linear proportionality on the initial protein monomer concentration, *m*_tot_ (Fig. 3c), providing quantitative evidence that the completion of the mature fibril aggregation reaction corresponds to the total protein monomer pool reaching the mature fibril state. These findings demonstrate that the mass concentration of mature fibrils at the end of the reaction is determined by the total protein monomer concentration, rather than the monomeric fraction available after the lipidic fibrils have formed. Taken together, our analysis indicates that the lipidic fibrils catalyze mature fibril formation and are on-pathway intermediates that are eventually consumed, with all protein in the system reaching the mature fibril state. This observation necessitates a kinetic model that accounts for the redistribution of protein mass from the metastable lipidic fibril to mature fibril form.

### Lipidic fibril dissolution is the rate-limiting step in mature fibril aggregation

To unify our mechanistic observations summarized in Fig. 3d, we constructed a global kinetic model to capture the full timecourse of the double sigmoid aggregation kinetics (Fig. 4a). This framework couples our established kinetic model for lipidic fibril aggregation with a description of the secondary nucleation-dominated aggregation of mature fibrils for the first and second growth phases, respectively. Crucially, the transition between these two regimes is mediated by heterogeneous primary nucleation. Finally, to account for the dissolution of lipidic fibrils, we introduce a coarse-grained description of the transfer of protein monomers from the lipidic to mature fibrils defined as a transfer rate, *r*_*T*_ . In this scenario, the lipidic and mature fibrils coexist and compete for a shared pool of protein monomers to elongate. The rapidly formed lipidic fibrils represent a metastable state in the structural land-scape with a critical concentration higher than that of the mature fibrils. Once the more thermodynamically stable mature fibrils are present, a net flux of monomers driven by the solubility differential is established because monomers dissociate from the lipidic fibril ends and are preferentially recruited by the elongating mature fibrils. The switch to a dissolution-controlled exchange between aggregated states ensures the eventual conversion of the entire protein pool to the mature fibril state, consistent with the dependence of ΔThT on the total protein concentration (Fig. 3c) and EM images (Fig. 1b-d).

**FIG. 4.**
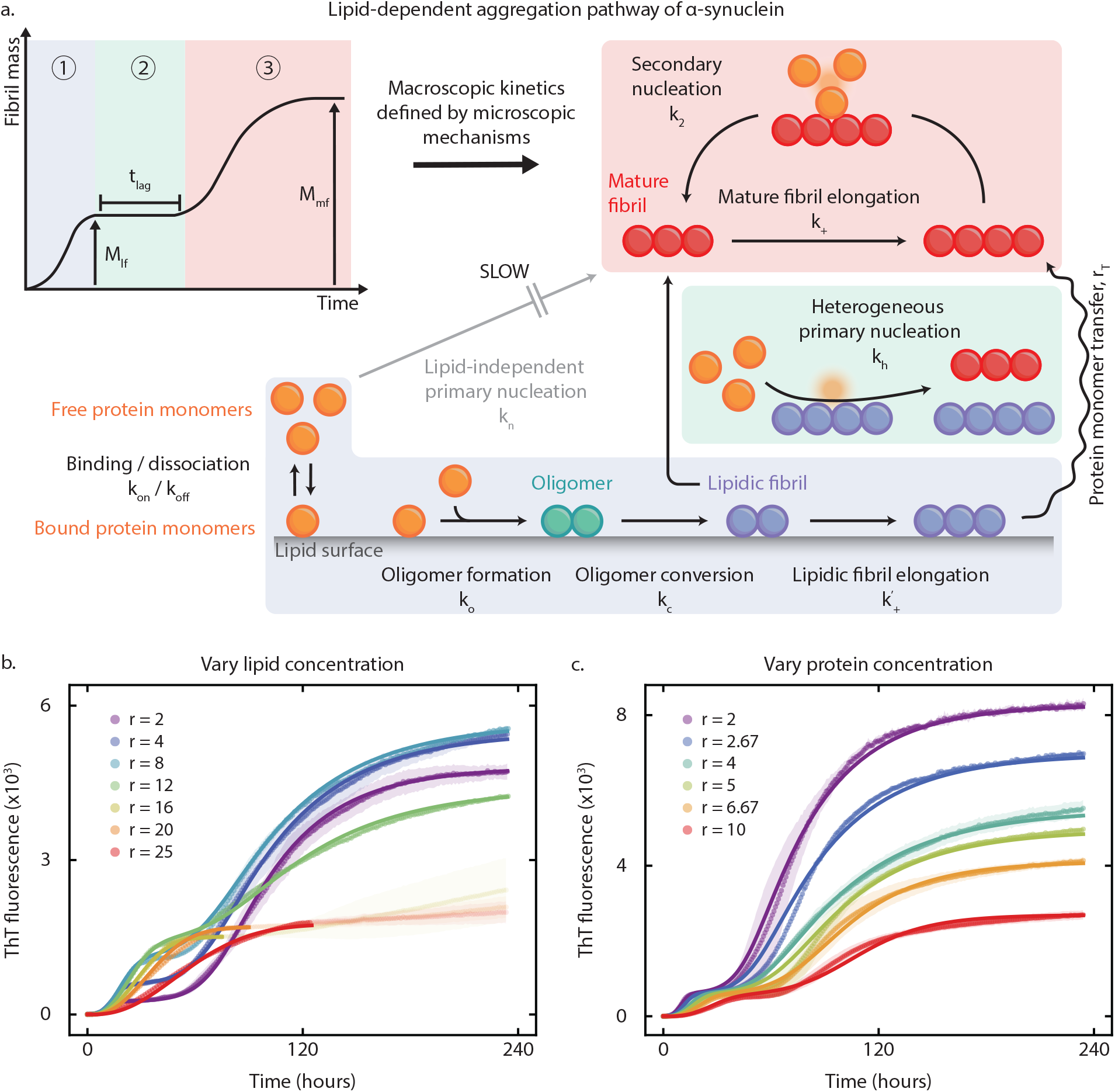
Kinetic analysis of *in vitro* lipid-dependent *α*-synuclein aggregation in polystyrene plate reveals the mechanistic pathway. **(a)** The lipid-dependent aggregation of *α*-synuclein to form mature fibrils consists of three distinct stages. (I) The rapid coaggregation of protein monomers and lipids forms lipidic fibrils, which, when present, enable (II) the formation of mature fibrils by a heterogeneous primary nucleation mechanism that controls the length of the intermediate plateau (*t*_lag_). (III) With mature fibrils present in the system, autocatalytic propagation is possible due to secondary nucleation, with the kinetics of the final stages of aggregation limited by lipidic fibril dissociation. Colored text labels define the different species relevant to the mechanistic pathway, and black text labels define microscopic reaction steps and the associated rate parameters. In the absence of the lipid-dependent pathway, mature fibril aggregation can proceed via the slower PS-induced primary nucleation. Experimental ThT fluorescence measurements of the time evolution of *in vitro* lipid-dependent *α*-synuclein aggregation, under varying initial **(b)** lipid and **(c)** *α*-synuclein monomer concentrations, where the experimental conditions are described in Fig. 2. Solid lines are best global fits to the self-consistent analytical solution describing lipidic fibril, Eq. (1), and mature fibril aggregation, Eq. (6). Aggregation curves for *r > α* are transparent in **(b)** as all protein monomers are consumed in the aggregation of lipidic fibrils in the first growth phase (Eq. (13)). The parameters describing lipidic fibril aggregation are taken from Fig. 2. **(b)** Fitted parameters: 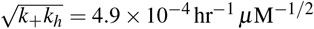 and 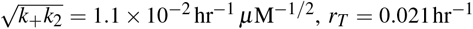 and *n*_*s*_ = 0.59 with constant *n*_*h*_ = 0. **(c)** Fitted parameters: 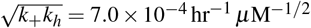 and 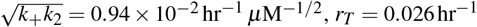 with constant *n*_*s*_ = 0 and *n*_*h*_ = 0. A summary of all fitting parameters can be found in Supplementary Material Table II.

If lipidic fibril dissolution is not included in the kinetic model, a systematic divergence between the model prediction and experimental data during the final approach to the steady state is observed (Supplementary Material Fig. S5). The full details of the kinetic model are provided in the Methods section.

We then fitted this model to the double-sigmoid ThT fluorescence measurements of lipid-dependent *α*-synuclein aggregation in the PS plates when either the initial lipid (Fig. 4b) or protein monomer concentration was varied (Fig. 4c). We used a single experimental readout of ThT fluorescence to monitor the concentration of both the lipidic and mature fibril species, which is possible because there is a clear separation of timescales over which these distinct fibrils form to produce the double sigmoid kinetic profile with the intermediate plateau between the growth phases. Furthermore, the two fibril species have different fluorescence quantum yields, which can be calculated using the predicted yield of lipidic fibrils after the first growth phase (Eq. (13)) and mature fibrils at the end of the aggregation reaction (Eq. (14)), with the measured ThT fluorescence in the kinetic experiments. The reaction order describing the dependence of heterogeneous primary nucleation on the concentration of protein monomer was set as *n*_*h*_ = 0 in our analysis, indicative of the nucleation being saturated with respect to the protein monomer contribution. As *r* approaches the critical point of *α*, less protein monomer remains after the aggregation of lipidic fibrils, transitioning the system to the sub-saturation regime, so *n*_*s*_ was instead treated as a fitting parameter in the analysis of the dataset when the initial concentration of lipids is varied. This leaves the reaction order *n*_*l*_ and the kinetic rates, 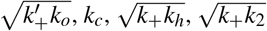, and *r*_*T*_, as fitting parameters in our analysis. The parameters associated with the second growth phase describe the rate of mature fibril elongation (*k*_+_), heterogeneous primary nucleation of mature fibrils (*k*_*h*_) and secondary nucleation (*k*_2_). The coupled rate parameters in our theoretical model (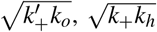 and *k*_+_*k*_2_) are quoted as 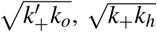 and 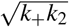 in this study, as these quantities are indicative of the timescales of lipidic and mature fibril propagation. These parameters are determined by globally fitting the normalized experimental data, segmented into first and second growth phases, to our theoretical model for lipidic (Eq. (1)) and mature fibril aggregation (Eq. (6)) respectively, as described in Supplementary Material Fig. S6. Using a secondary nucleation-dominated kinetic model to describe the propagation of mature fibrils was supported by our analysis of lipid-independent *α*-synuclein aggregation data (Supplementary Material Fig. S7). It has been previously demonstrated that secondary nucleation does not feature in the mechanistic landscape of lipid-independent *α*-synuclein aggregation under the experimental conditions used in this study in PEGylated plates^18^. However, as PS facilitates both the homogeneous and heterogeneous primary nucleation of mature fibrils (no double sigmoid aggregation profiles observed in PEGylated plates), secondary nucleation could also be induced by the PS via similar mechanisms of surface-protein interactions.

### Ostwald’s Rule of Stages and the free energy landscape of *α*-synuclein aggregation

Our observation that the lipid-dependent aggregation of *α*-synuclein first forms the metastable lipidic fibril intermediate that, once mature fibrils are present in the system, undergoes a solubility-driven dissolution transferring protein monomers from the lipidic to mature fibril state confirms that the lipid-dependent aggregation of *α*-synuclein follows Ostwald’s Rule of Stages in our experimental assay (Fig. 5). Therefore, the double sigmoid kinetic profile represents the sequential descent through the free energy landscape, where *α*-synuclein transitions through a series of increasingly more stable states. Elucidating the mechanistic landscape of aggregation reveals a clear hierarchy of stability in which less stable monomers initially form the kinetically favorable lipidic fibrils before the thermodynamically more stable mature fibrils form (*G*_*m*_ *> G*_*l f*_ *> G*_*m f*_ ). Once the more stable mature fibrils are present in the system, the thermodynamic gradient due to a negative Δ*G*_*l f,m f*_, establishes a chemical potential gradient (*µ*_*l f*_ *> µ*_*m f*_ ) between the two fibril species, which drives the dissolution of lipidic fibrils. The existence of the intermediate plateau and the protracted growth tail in the aggregation data are therefore fundamental consequences of the system navigating a multi-step energy landscape. Furthermore, the free energy landscape rationalizes why the lipid-dependent aggregation pathway is kinetically favorable. Despite the fact that the PS surface provides a heterogeneous nucleation pathway for mature fibrils, the PS does not incorporate into the fibril structure to form an intermediate fibril structure as the lipid surface does, so the aggregation cannot proceed through a sequential formation of increasingly stable states. To characterize the kinetic accessibility of the free energy landscape, we calculated the activation energies associated with the protein state transitions in the lipid-dependent *α*-synuclein aggregation pathway using the kinetic rate parameters quantified by our kinetic framework in Fig. 4. The formation of lipidic fibrils (described by the combined rate parameter *µ* Eq. (5)) is significantly more kinetically accessible compared to PS-induced primary nucleation of mature fibrils (described by the combined rate parameter *λ* where *k*_*h*_ = 0, Eq. (8)). We find that the presence of the lipidic fibril surface lowers the barrier to mature fibril nucleation by Δ*E*_*a,m f*_ *∼* 25 kJmol^*−*1^ relative to the PS-induced primary nucleation in the lipid-independent pathway, where 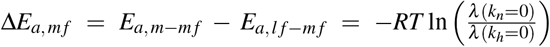 (See Eq. (8) for definition of *λ* ), and explains the acceleration of mature fibril aggregation in the lipid-dependent pathway. As we use PS to accelerate lipid-independent aggregation, Δ*E*_*a,m f*_ would increase in the absence of the heterogeneous nucleation pathway provided by the PS surface, further highlighting the critical role lipidic fibrils play in accelerating mature fibril aggregation.

**FIG. 5.**
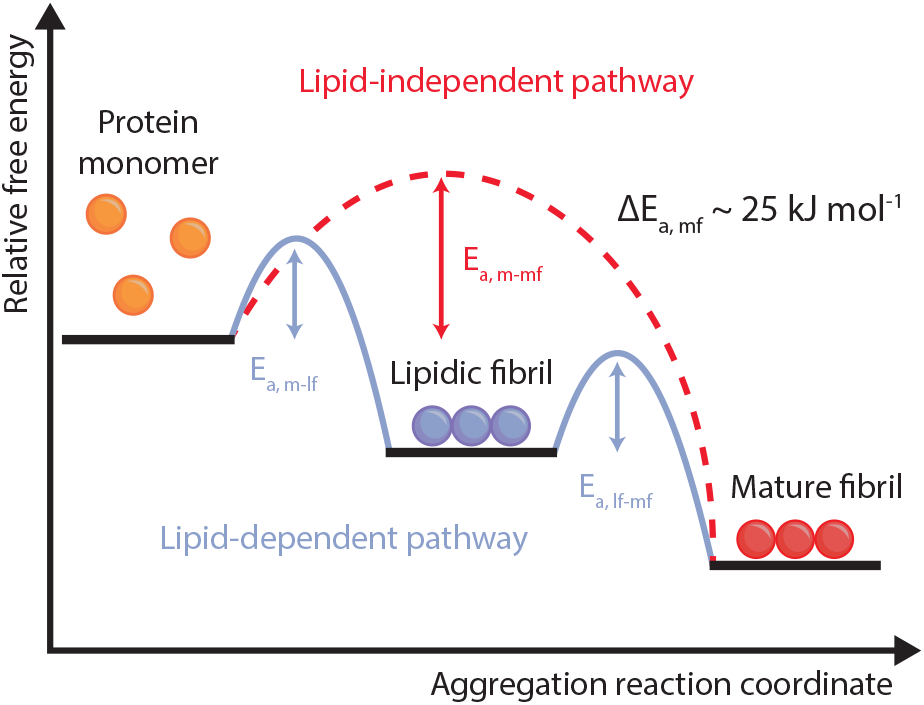
Lipid-dependent aggregation of mature *α*-synuclein fibrils follows Ostwald’s Rule of Stages and lowers the activation energy barrier for mature fibril nucleation. Schematic energy landscape for *α*-synuclein aggregation via lipid-independent and lipid-dependent pathways. As the presence of mature fibrils induces the dissolution of lipidic fibrils, the mature fibril state must be thermo-dynamically more stable than the lipidic fibrils. Therefore, the lipid-dependent aggregation pathway follows Ostwald’s Rule of Stages as fibril species of increasing stability are sequentially produced in the aggregation reaction. To quantify the kinetic accessibility of the lipidic and mature fibril states, we calculate the difference in activation energies associated with the aggregation pathways using the kinetic rate parameters identified in Fig. 4. The lipidic fibril surface lowers the barrier to mature fibril nucleation by *E*_*a, m−m f*_ *−E*_*a, l f −m f*_ *∼* 25 kJmol^*−*1^ relative to the PS surface. Relative state minima illustrate increasing thermodynamic stability from monomer to mature fibril, and barrier heights represent apparent activation energies (*E*_*a*_).

### Ambroxol specifically modulates double sigmoid kinetics by targeting the lipid-dependent pathway

To further validate our proposed mechanistic landscape, we utilized the small molecule ambroxol (ABX) as a selective probe to inhibit the coaggregation step in the lipid-dependent pathway. We have previously demonstrated that ABX specifically inhibits the coaggregation of *α*-synuclein with lipids, while lipid-independent aggregation is not affected^36^. The inhibitory signature of ABX, with a dose-dependent reduction in both the lipidic fibril mass yield and the oligomer formation rate, was fully preserved in our PS plate assay. Increasing the concentration of ABX progressively attenuated the first growth phase, reflecting a decrease in the concentration of the lipidic fibrils, and delayed the onset of the second growth phase (Fig. 6a). Crucially, at sufficiently high concentrations of ABX where the formation of lipidic fibrils was completely inhibited, the double sigmoid profile is no longer present. In this regime, the kinetic profile collapsed onto the slow, single sigmoid kinetics identical to the lipid-independent aggregation profile (Fig. 6a). The convergence of the kinetic profiles demonstrates that without the lipidic fibril scaffold, the system loses the catalytic surface required for heterogeneous primary nucleation and mature fibril nuclei can only be formed by PS-induced primary nucleation. This is evidenced by the half-time of aggregation (Fig. 6b), where the addition of lipids decreases the half-time by *∼* 50%, which is then attenuated in an ABX concentration-dependent fashion. To confirm that this modulation is specific to the chemical properties of ABX rather than a generic small molecule interference, we used conduritol B epoxide (CBE) at a concentration where ABX achieves complete inhibition of lipidic fibril aggregation. CBE failed to produce any significant inhibitory effect on either growth phase, reinforcing that the observed kinetic modulation is a specific consequence of ABX-lipid-protein interactions (Fig. 6b and Supplementary Material Fig. S8). This selective inhibition of the lipid-dependent pathway of aggregation confirms that the accelerated formation of mature fibrils is strictly dependent on the presence of the metastable lipidic fibril intermediate.

**FIG. 6.**
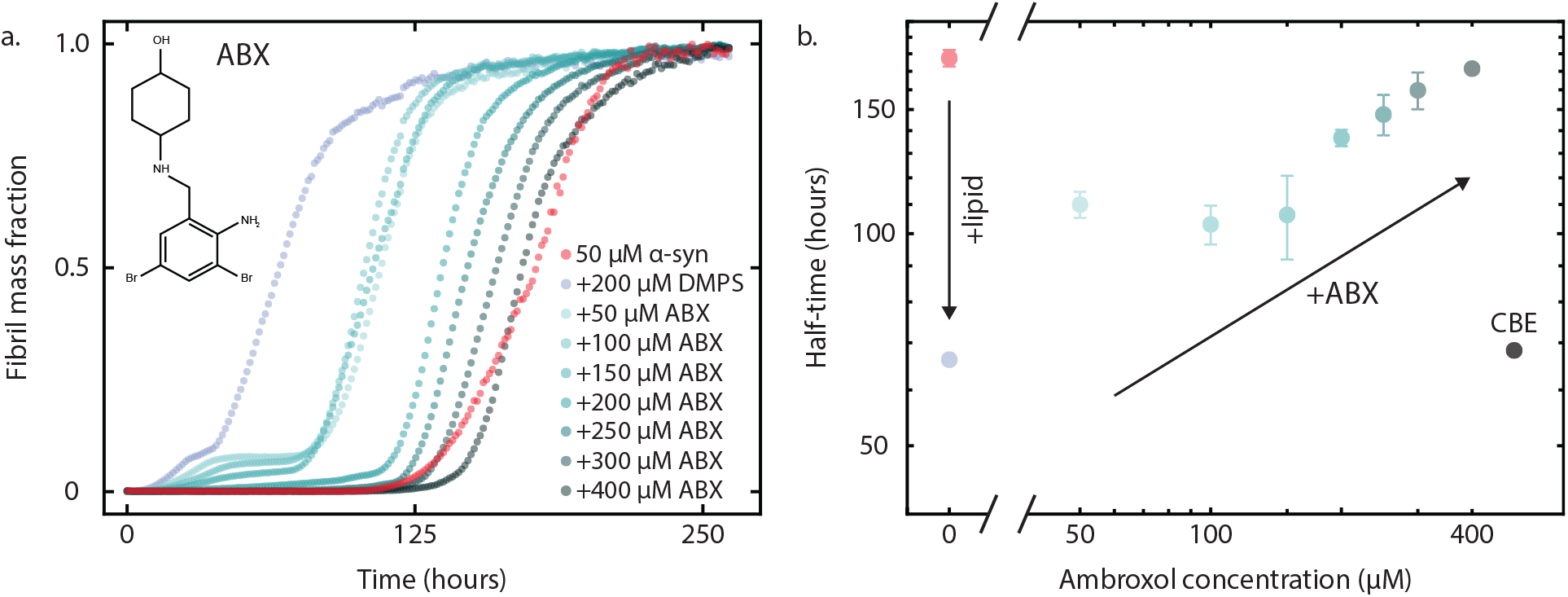
Selective inhibition of lipidic fibril formation attenuates the accelerated lipid-dependent *α*-synuclein aggregation pathway. **(a)** Experimental aggregation curves measured with ThT fluorescence with constant *m*_tot_ = 50 *µ*M and *L*_tot_ = 200 *µ*M (no lipid in red curve) and varying concentrations of ambroxol (50 *−* 400 *µ*M). **(b)** Half-times of mature fibril aggregation (when fibril mass fraction reaches 0.5 in panel (a)) highlight the acceleration upon the addition of DMPS lipids that is attenuated in a concentration-dependent fashion by ambroxol. The inhibition of mature fibrils is not a generic small molecule effect, as the addition of 400 *µ*M of CBE does not change the half-time (kinetic curve shown in Supplementary Material Fig. S8).

An alternative mechanistic landscape for the lipid-dependent aggregation pathway is that the DMPS lipids bind directly to the PS surface and thus passivate the catalytic activity of the PS to induce mature fibril formation. In this scenario, the rapid formation of lipidic fibrils would remove the lipids bound to the PS surface, increasing the catalytic surface area available for PS-induced aggregation, which would also predict a shorter *t*_lag_ with increasing lipidic fibril yield (by increasing *L*_tot_). However, the inhibitory effect of ABX on the aggregation kinetics provides quantitative evidence disproving this scenario, as when the formation of lipidic fibrils is inhibited by ABX, the accelerated aggregation of mature fibrils is also attenuated despite the fact that lipids are still present in the system. Our seeding experiments also support our proposed interpretation of the mechanistic landscape, as the presence of increasing concentrations of lipidic fibrils, not lipids, increases the rate of heterogeneous primary nucleation of mature fibrils (Fig. 3b). Furthermore, as both the PS surface in the microwell plate and DMPS lipid are negatively charged, direct interaction and binding is unlikely.

## III. CONCLUSIONS

We have developed a kinetic framework to rationalize the double-sigmoid kinetics of *α*-synuclein aggregation to form mature fibrils in the presence of lipids in PS plates, across a broad range of protein and lipid concentrations. We have revealed that lipidic fibrils serve as a catalytic surface that accelerates the aggregation of mature fibrils through a heterogeneous primary nucleation mechanism. Crucially, the double-sigmoid kinetic profile is not unique to the lipid-dependent aggregation in the PS plate, as it has been observed when temperature is increased to induce the shedding of lipids from the lipidic fibrils^4^. Lipid-induced aggregation has been observed for other amyloid-forming systems such as islet amyloid polypeptide and amyloid-*β* ^57,58^. Furthermore, our analysis confirms that the relatively slow timescale of aggregation of mature fibrils reaching steady state is due to fibril elongation becoming limited by the rate at which protein monomers dissociate from the metastable lipidic fibrils. The dissolution of lipidic fibrils in the presence of the thermodynamically more stable mature fibrils confirms that the fibril species form in order of stability, following Ostwald’s Rule of Stages. Our findings, coupled with the fact that mutations in the gene GBA encoding the enzyme Glucocerebrosidase (GCase) are together the most important genetic risk factor for developing PD and are associated with alterations in the levels and properties of GCase’s lipid substrates and products^41,62^, suggest that the dysregulation of lipids could be a critical initiator of *α*-synuclein aggregation to form mature amyloid fibrils. Pathological dysregulation of lipid levels could first enable the kinetically favorable aggregation of lipidic fibrils, whose presence then accelerates the aggregation of mature fibrils and the subsequent cytotoxic effects associated with the transformation of functional *α*-synuclein monomers to the amyloid state.

Furthermore, our findings provide a possible explanation as to why no lipids are found in *ex vivo* amyloid fibrils, as they are only part of the metastable intermediate fibrils. By employing DMPS as a model membrane system, selected because phosphatidylserine is a major component of biological membranes and has been shown to yield highly reproducible lipidic fibril aggregation kinetics, enabled the development of a generalizable quantitative foundation for studying surface-catalyzed amyloid fibril aggregation in biologically relevant environments. Although our experimental assay utilizes PS plates, the role of the PS surface is primarily to enable a direct, quantitative comparison of lipid-dependent and lipid-independent aggregation pathways given the slow homogeneous primary nucleation of *α*-synuclein, rather than act as the central driver of the mechanistic landscape of aggregation.

Our proposed mechanistic pathway of lipid-dependent mature fibril aggregation is further supported by the kinetic effects of selectively inhibiting the formation of the lipidic fibrils by ABX. Removing the heterogeneous primary nucleation of mature fibrils facilitated by the lipidic fibrils collapses the aggregation kinetics back to the slow lipid-independent kinetic profile. Unlike amyloid inhibition strategies that aim to stabilize protein monomers or disaggregate mature fibrils while not accounting for heterogeneous protein interactions that could still induce aggregation, an inhibitor such as ABX could act upstream of mature fibril aggregation by stopping the formation of the lipidic fibril catalyst. By inhibiting the accelerated lipid-dependent aggregation pathway, ABX could delay the associated cytotoxic effects of amyloid aggregation. Despite ABX attenuating the accelerated aggregation facilitated by lipidic fibrils, mature fibrils can still form via the lipid-independent pathway in our *in vitro* assay. Therefore, a possible therapeutic strategy could involve inhibiting both pathways of mature fibril aggregation described in this study, where a combination therapy that also includes an inhibitor of the lipid-independent pathway could achieve higher efficacy. Taken together, these findings provide a quantitative basis for understanding how heterogeneous surfaces can fundamentally alter the energy landscape and kinetics of *α*synuclein aggregation and provide a foundation for designing novel strategies for the targeted pharmacological disruption of lipid-accelerated amyloid aggregation.

This work was funded by ETH Zurich (AS, TCTM), the Swiss National Science Foundation (grant no 219703 to TCTM) and the Novo Nordisk Foundation (Grant number: NNF20OC0059417 to CG) (JED, NTC, CG)

## AUTHOR CONTRIBUTIONS

A.S., J.E.D., C.G., and T.C.T.M. designed research; A.S., J.E.D., N.B., N.T.C., C.G., and T.C.T.M. performed research; A.S., J.E.D., C.G., and T.C.T.M. contributed new reagents/analytic tools; A.S., J.E.D., N.B., N.T.C., C.G., and T.C.T.M. analyzed data; and A.S., J.E.D., N.B., N.T.C., C.G., and T.C.T.M. wrote the paper.

## COMPETING INTERESTS

The authors declare no competing interests.

## METHODS

### Integrated rate laws for lipidic fibril aggregation

Here, we provide explicit formulas for the integrated rate laws describing lipidic fibril aggregation^27,30^. The analytical expression for the lipidic fibril mass concentration can be written as

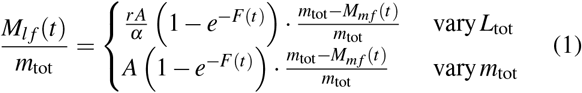

where *r* = *L*_tot_*/m*_tot_ is the lipid-to-monomer ratio, *α* is the average number of lipid units bound by a single protein monomer in lipidic fibril form and

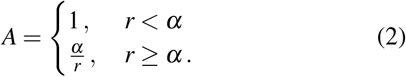

The function *F*(*t*) depends on the dominant mechanism of lipidic fibril nucleation. For two-step primary nucleation

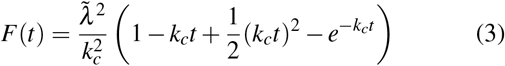

where

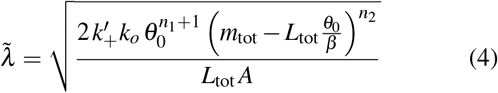

is an effective rate constant of lipidic fibril proliferation through oligomer formation and lipidic fibril elongation. Here, 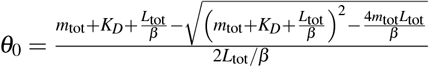 where *K*_*D*_ = *k*_off_*/k*_on_ is the equilibrium dissociation constant of protein-lipid binding and *β* is the average number of lipid units bound by a single protein monomer. We define *k*_*o*_ as the rate of oligomer formation, with reaction orders *n*_1_ and *n*_2_ that describe the dependence of oligomer formation on bound and free protein monomers, respectively. The rate of conversion of oligomers to lipidic fibrils is described by *k*_*c*_ and the rate of lipidic fibril elongation is 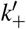. The final term in Eq. (1), (*m*_tot_ *− M*_*m f*_ (*t*))*/m*_tot_, couples lipidic fibril mass to the growing mature fibril population and accounts for the dissolution of lipidic fibrils as mature fibrils aggregate: as *M*_*m f*_*→ m*_tot_, the lipidic fibril mass is driven to zero, redistributing protein into the mature fibril state. The rate of this redistribution is governed by the transfer parameter *r*_*T*_ . Because we rely on a single experimental readout, ThT fluorescence, which cannot separately resolve the two fibril species, we do not measure the dynamics of lipidic fibril dissolution directly. Instead, *r*_*T*_ is identified indirectly from the shape of the late-time kinetics: the rate-limiting dissolution manifests as the characteristically slow approach to the final ThT plateau, which a model without the dissolution term fails to capture.

We define an effective rate of lipidic fibril proliferation as

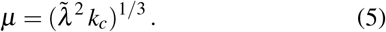

Examining the half-time scaling behavior of lipidic fibril mass concentration using Eq. (1) reveals the scaling exponents and their dependence on the relevant reaction orders^27,30^. These scaling results were used to identify the reaction orders, *n*_1_ and *n*_2_, in this study. For *r < α*
where *γ*_*p*_ and *γ*_*L*_ describe the dependence of *t*_1*/*2_ on *m*_tot_ and *L*_tot_ being varied, respectively. For *r > α*

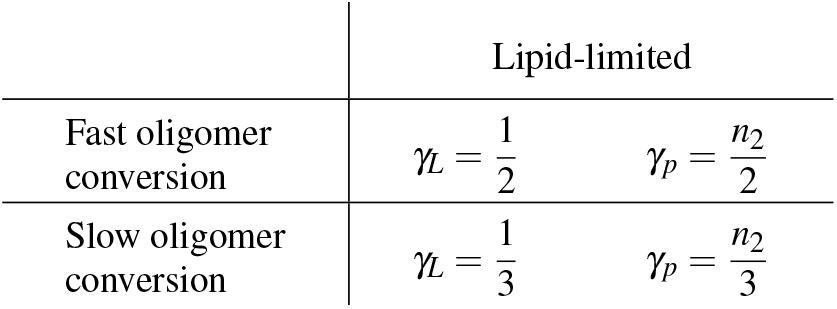

|  | Protein-limited |  |
|---|---|---|
| Fast oligomer conversion | $\gamma_L = \frac{n_1 + n_2 + 1}{2}$ | $\gamma_p = \frac{n_1 + n_2}{2}$ |
| Slow oligomer conversion | $\gamma_L = \frac{n_1 + n_2 + 1}{3}$ | $\gamma_p = \frac{n_1 + n_2}{3}$ |

### Integrated rate laws for mature fibril aggregation

The analytical expression for the mature fibril mass concentration can be written as

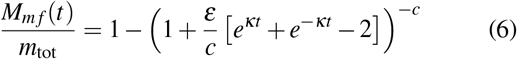

where

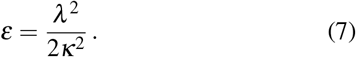

The *λ* term describes the effective rate of primary nucleation of mature fibrils, through both PS-induced and heterogeneous mechanisms, and is defined as

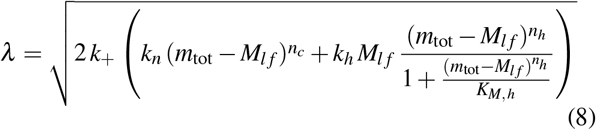

where *k*_+_ is the rate of elongation of mature fibrils, *k*_*n*_ is the rate of PS-induced primary nucleation with associated reaction order *n*_*c*_ defining the dependence of this process on the protein monomer concentration remaining after lipidic fibril aggregation. Similarly, *k*_*h*_ is the rate of heterogeneous primary nucleation of mature fibrils with associated reaction order *n*_*h*_. As heterogeneous primary nucleation is inherently a multistep process, where protein monomers must first bind to the surface of the lipidic fibrils before they can form a stable mature fibril nucleus that then dissociates from the lipidic fibril surface, saturation can occur. The saturation effects can be determined by a single constant, *K*_*M,h*_, which plays a role analogous to the Michaelis constant in enzymology and represents a threshold concentration above which the protein monomer dependence of the heterogeneous primary nucleation process becomes saturated^63^.

The *κ* term describes the effective rate of secondary nucleation of mature fibrils and is defined as

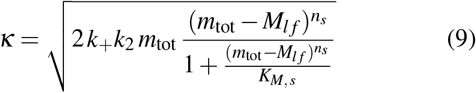

where *k*_2_ is the rate of secondary nucleation of mature fibrils with associated reaction order *n*_*s*_ defining the dependence of this process on the free protein monomer concentration remaining after lipidic fibril aggregation. Similarly, we have accounted for the saturation of the secondary nucleation process (when all catalytic sites on the mature fibril sites are occupied) with the constant *K*_*M,s*_.

Finally, by accounting for the rate-limiting timescale of lipidic fibril dissolution to enable protein monomer transfer to the elongating mature fibrils, *c* is defined as

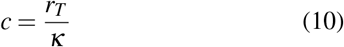

where *r*_*T*_ is the transfer rate of protein monomers from lipidic to mature fibrils. Therefore, 1*/c* defines a Damköhler number for our system as the ratio of the effective rate of mature fibril aggregation (*κ*) and the transport phenomena rate occurring in the system (*r*_*T*_ ), which is the 1*/c* term in our mathematical model, Eq. (6)). In theoretical descriptions of homogeneous aggregation kinetics, 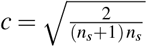.

In the presence of lipidic fibrils, the reaction flux of PS-induced primary nucleation is negligible relative to the heterogeneous primary nucleation flux. In our analysis of experimental aggregation curves in Fig. 4, we neglect PS-induced primary nucleation for this reason. Furthermore, as all kinetic curves where a second growth phase is consistently observed in Fig. 4 are in the saturated regime for the protein monomer dependency of heterogeneous primary nucleation, we further simplify Eq. (8) to

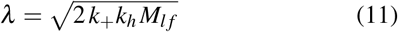

in our fitting analysis to minimize the number of fitting parameters required to interpret the experimental data.

### Steady-state behavior

At the end of the first growth phase, the yield of lipidic fibrils is given by

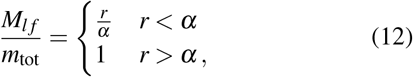

where the dependence on the second growth phase being feasible, requiring protein monomers to remain in the system after the lipidic fibril aggregation is complete, on *r* is clear. The steady-state lipidic fibril mass concentration is defined, from Eq. (1), as

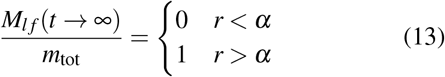

and for the mature fibril mass concentration, from Eq. 6 we find

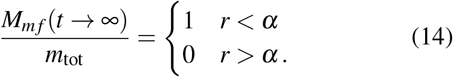

### Combined model and fitting experimental data

As the single experimental readout of ThT fluorescence is used to track the mass concentrations of both fibril species, we rely upon the different quantum yields of fluorescence from each fibril in our fitting analysis of the data. The measured ThT fluorescence after the first and second growth phase is divided by the predicted steady-state concentrations of lipidic and mature fibrils (Eq. (13) and Eq. (14)) to quantify the quantum yield. With the quantum yields, we combined Eq. (1) and Eq. (6) to produce

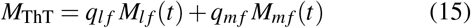

where *q*_*l f*_ and *q*_*m f*_ are the quantum yields of lipidic and mature fibrils, respectively. *M*_ThT_ is the output quantity plotted with the raw ThT signal in Fig. 4. To improve the quality of the global fitting analysis, the data were segmented into the first growth phase and from the intermediate plateau to the end of the second growth phase, which were normalized before fitting with Eq. (1) and Eq. (6).

## Notes

### Competing Interest Statement

The authors have declared no competing interest.

## References

1 W. Ostwald, “Studien über die bildung und umwandlung fester körper: 1. abhandlung: Übersättigung und Überkaltung,” Zeitschrift für Physikalische Chemie 22U, 289–330 (1897).

2 J. L. Wray and F. Daniels, “Precipitation of calcite and aragonite,” Journal of the American Chemical Society 79, 2031–2034 (1957).

3 A. Levin, T. O. Mason, L. Adler-Abramovich, A. K. Buell, G. Meisl, C. Galvagnion, Y. Bram, S. A. Stratford, C. M. Dobson, T. P. J. Knowles, and E. Gazit, “Ostwald’s rule of stages governs structural transitions and morphology of dipeptide supramolecular polymers,” Nature Communications 5 (2014), 10.1038/ncomms6219.

4 J. W. P. Brown, G. Meisl, T. P. J. Knowles, A. K. Buell, C. M. Dobson, and C. Galvagnion, “Kinetic barriers to α-synuclein protofilament formation and conversion into mature fibrils,” Chemical Communications 54, 7854–7857 (2018).

5 C. M. Dobson, “Protein folding and misfolding,” Nature 426, 884–890 (2003).

6 F. Chiti and C. M. Dobson, “Protein misfolding, functional amyloid, and human disease,” Annual Review of Biochemistry 75, 333–366 (2006).

7 T. P. J. Knowles, M. Vendruscolo, and C. M. Dobson, “The amyloid state and its association with protein misfolding diseases,” Nature Reviews Molecular Cell Biology 15, 384–396 (2014).

8 F. Chiti and C. M. Dobson, “Protein misfolding, amyloid formation, and human disease: a summary of progress over the last decade,” Annual Review of Biochemistry 86, 27–68 (2017).

9 M. G. Spillantini, M. L. Schmidt, V. M.-Y. Lee, J. Q. Trojanowski, R. Jakes, and M. Goedert, “α-synuclein in Lewy bodies,” Nature 388, 839–840 (1997).

10 M. Hashimoto and E. Masliah, “Alpha-synuclein in Lewy body disease and Alzheimer’s disease,” Brain Pathology 9, 707–720 (1999).

11 A. Bellucci, M. Zaltieri, L. Navarria, J. Grigoletto, C. Missale, and P. Spano, “From α-synuclein to synaptic dysfunctions: New insights into the pathophysiology of Parkinson’s disease,” Brain Research 1476, 183–202 (2012).

12 M. G. Spillantini and M. Goedert, “The α-synucleinopathies: Parkinson’s disease, dementia with Lewy bodies, and multiple system atrophy,” Annals of the New York Academy of Sciences 920, 16–27 (2000).

13 I. Gureviciene, K. Gurevicius, and H. Tanila, “Role of α-synuclein in synaptic glutamate release,” Neurobiology of Disease 28, 83–89 (2007).

14 G. Fusco, A. De Simone, T. Gopinath, V. Vostrikov, M. Vendruscolo, C. M. Dobson, and G. Veglia, “Direct observation of the three regions in α-synuclein that determine its membrane-bound behaviour,” Nature Communications 5, 1–8 (2014).

15 K. Makasewicz, S. Wennmalm, B. Stenqvist, M. Fornasier, A. Andersson, P. Jönsson, S. Linse, and E. Sparr, “Cooperativity of α-synuclein binding to lipid membranes,” ACS Chemical Neuroscience 12, 2099–2109 (2021).

16 J. Burré, “The synaptic function of α-synuclein,” Journal of Parkinson’s Disease 5, 699–713 (2015).

17 A. L. Fink, “The aggregation and fibrillation of α-synuclein,” Accounts of Chemical Research 39, 628–634 (2006).

18 A. K. Buell, C. Galvagnion, R. Gaspar, E. Sparr, M. Vendruscolo, T. P. J. Knowles, S. Linse, and C. M. Dobson, “Solution conditions determine the relative importance of nucleation and growth processes in α-synuclein aggregation,” Proceedings of the National Academy of Sciences 111, 7671–7676 (2014).

19 C. K. Xu, G. Meisl, E. Andrzejewska, G. Krainer, A. J. Dear, M. C. Cruz, S. Turi, R. Jacquat, W. E. Arter, M. Vendruscolo, et al., “α-synuclein oligomers form by secondary nucleation,” bioRxiv, 2023–05 (2023).

20 C. K. Xu, G. Meisl, E. A. Andrzejewska, G. Krainer, A. J. Dear, M. Castellana-Cruz, S. Turi, I. A. Edu, G. Vivacqua, R. P. B. Jacquat, W. E. Arter, M. G. Spillantini, M. Vendruscolo, S. Linse, and T. P. J. Knowles, “-synuclein oligomers form by secondary nucleation,” Nature Communications 15 (2024), 10.1038/s41467-024-50692-4.

21 M. Zhu and A. L. Fink, “Lipid binding inhibits α-synuclein fibril formation,” Journal of Biological Chemistry 278, 16873–16877 (2003).

22 Z. Martinez, M. Zhu, S. Han, and A. L. Fink, “Gm1 specifically interacts with α-synuclein and inhibits fibrillation,” Biochemistry 46, 1868–1877 (2007).

23 L. Giehm, D. I. Svergun, D. E. Otzen, and B. Vestergaard, “Low-resolution structure of a vesicle disrupting α-synuclein oligomer that accumulates during fibrillation,” Proceedings of the National Academy of Sciences 108, 3246–3251 (2011).

24 C. Galvagnion, A. K. Buell, G. Meisl, T. C. T. Michaels, M. Vendruscolo, T. P. J. Knowles, and C. M. Dobson, “Lipid vesicles trigger α-synuclein aggregation by stimulating primary nucleation,” Nature Chemical Biology 11, 229 (2015).

25 P. Flagmeier, G. Meisl, M. Vendruscolo, T. P. J. Knowles, C. M. Dobson, A. K. Buell, and C. Galvagnion, “Mutations associated with familial Parkinson’s disease alter the initiation and amplification steps of α-synuclein aggregation,” Proceedings of the National Academy of Sciences 113, 10328–10333 (2016).

26 C. Galvagnion, A. Barclay, K. Makasewicz, F. R. Marlet, M. Moulin, J. M. Devos, S. Linse, A. Martel, L. Porcar, E. Sparr, M. C. Pedersen, F. Roosen-Runge, L. Arleth, and A. K. Buell, “Structural characterisation of α-synuclein-membrane interactions and the resulting aggregation using small angle scattering,” Physical Chemistry Chemical Physics 26, 10998–11013 (2024).

27 A. Stevenson, R. Staats, A. J. Dear, D. Voderholzer, J. E. Dreier, G. Meisl, R. Guido, T. P. J. Knowles, C. Galvagnion, A. K. Buell, M. Vendruscolo, and T. C. T. Michaels, “Global kinetic model of lipid-induced α-synuclein aggregation and its inhibition by small molecules,” Proceedings of the National Academy of Sciences 122 (2025), 10.1073/pnas.2422427122.

28 C. Galvagnion, J. W. P. Brown, M. M. Ouberai, P. Flagmeier, M. Vendruscolo, A. K. Buell, E. Sparr, and C. M. Dobson, “Chemical properties of lipids strongly affect the kinetics of the membrane-induced aggregation of α-synuclein,” Proceedings of the National Academy of Sciences 113, 7065–7070 (2016).

29 J. Ramirez, S. X. Pancoe, E. Rhoades, and E. J. Petersson, “The effects of lipids on α-synuclein aggregation in vitro,” Biomolecules 13, 1476 (2023).

30 A. Stevenson, D. Voderholzer, and T. C. T. Michaels, “Self-consistent analytical solutions to the kinetics of lipid-induced protein aggregation,” The Journal of Chemical Physics 163 (2025), 10.1063/5.0279601.

31 E. Hellstrand, A. Nowacka, D. Topgaard, S. Linse, and E. Sparr, “Membrane lipid co-aggregation with α-synuclein fibrils,” PLoS ONE 8, e77235 (2013).

32 B. Frieg, L. Antonschmidt, C. Dienemann, J. A. Geraets, E. E. Najbauer, D. Matthes, B. L. de Groot, L. B. Andreas, S. Becker, C. Griesinger, et al., “The 3d structure of lipidic fibrils of α-synuclein,” Nature Communications 13, 6810 (2022).

33 M. L. Choi, A. Chappard, B. P. Singh, C. Maclachlan, M. Rodrigues, E. I. Fedotova, A. V. Berezhnov, S. De, C. J. Peddie, D. Athauda, G. S. Virdi, W. Zhang, J. R. Evans, A. I. Wernick, Z. S. Zanjani, P. R. Angelova, N. Esteras, A. Y. Vinokurov, K. Morris, K. Jeacock, L. Tosatto, D. Little, P. Gissen, D. J. Clarke, T. Kunath, L. Collinson, D. Klenerman, A. Y. Abramov, M. H. Horrocks, and S. Gandhi, “Pathological structural conversion of α-synuclein at the mitochondria induces neuronal toxicity,” Nature Neuroscience 25, 1134–1148 (2022).

34 A. D. Stephens, A. F. Villegas, C. W. Chung, O. Vanderpoorten, D. Pinotsi, I. Mela, E. Ward, T. M. McCoy, R. Cubitt, A. F. Routh, C. F. Kaminski, and G. S. Kaminski Schierle, “α-synuclein fibril and synaptic vesicle interactions lead to vesicle destruction and increased lipid-associated fibril uptake into ipsc-derived neurons,” Communications Biology 6 (2023), 10.1038/s42003-023-04884-1.

35 C. Galvagnion, D. Topgaard, K. Makasewicz, A. K. Buell, S. Linse, E. Sparr, and C. M. Dobson, “Lipid dynamics and phase transition within α-synuclein amyloid fibrils,” The Journal of Physical Chemistry Letters 10, 7872–7877 (2019).

36 J. E. Dreier, A. Stevenson, E. Carles, K. Schott, T. C. T. Michaels, and C. Galvagnion, “Ambroxol displaces -synuclein from the membrane and inhibits the formation of early protein–lipid coaggregates,” Chemical Science 17, 354–363 (2026).

37 D. Eliezer, E. Kutluay, R. Bussell, and G. Browne, “Conformational properties of -synuclein in its free and lipid-associated states 1 1edited by p. e. wright,” Journal of Molecular Biology 307, 1061–1073 (2001).

38 Y. Yang, Y. Shi, M. Schweighauser, X. Zhang, A. Kotecha, A. G. Murzin, H. J. Garringer, P. W. Cullinane, Y. Saito, T. Foroud, T. T. Warner, K. Hasegawa, R. Vidal, S. Murayama, T. Revesz, B. Ghetti, M. Hasegawa, T. Lashley, S. H. W. Scheres, and M. Goedert, “Structures of -synuclein filaments from human brains with lewy pathology,” Nature 610, 791–795 (2022).

39 R. Vácha, S. Linse, and M. Lund, “Surface effects on aggregation kinetics of amyloidogenic peptides,” Journal of the American Chemical Society 136, 11776–11782 (2014).

40 R. Gaspar, G. Meisl, A. K. Buell, L. Young, C. F. Kaminski, T. P. J. Knowles, E. Sparr, and S. Linse, “Secondary nucleation of monomers on fibril surface dominates α-synuclein aggregation and provides autocatalytic amyloid amplification,” Quarterly Reviews of Biophysics 50, e6 (2017).

41 C. Galvagnion, F. R. Marlet, S. Cerri, A. H. V. Schapira, F. Blandini, and D. A. Di Monte, “Sphingolipid changes in parkinson l444p gba mutation fibroblasts promote α-synuclein aggregation,” Brain 145, 1038–1051 (2022).

42 M. Grey, C. J. Dunning, R. Gaspar, C. Grey, P. Brundin, E. Sparr, and S. Linse, “Acceleration of -synuclein aggregation by exosomes,” Journal of Biological Chemistry 290, 2969–2982 (2015).

43 H. Mohammad-Beigi, M. Zanganeh, C. Scavenius, H. Eskandari, A. Farzadfard, S. A. Shojaosadati, J. J. Enghild, D. E. Otzen, A. K. Buell, and D. S. Sutherland, “A protein corona modulates interactions of -synuclein with nanoparticles and alters the rates of the microscopic steps of amyloid formation,” ACS Nano 16, 1102–1118 (2022).

44 Z. Liu, A. Sokratian, A. M. Duda, E. Xu, C. Stanhope, A. Fu, S. Strader, H. Li, Y. Yuan, B. G. Bobay, J. Sipe, K. Bai, I. Lundgaard, N. Liu, B. Hernandez, C. Bowes Rickman, S. E. Miller, and A. B. West, “Anionic nanoplastic contaminants promote parkinson’s disease–associated α-synuclein aggregation,” Science Advances 9 (2023), 10.1126/sci-adv.adi8716.

45 A. Mishra, T. W. Golbek, A. B. Thomassen, L. Zuzic, L. Schmüser, K. H. Saeed, F. Madzharova, J. Nielsen, B. Schiøtt, D. E. Otzen, and T. Weidner, “Pathological folding of -synuclein on polystyrene nanoplastic revealed by sum frequency scattering and 2d infrared spectroscopy,” The Journal of Physical Chemistry Letters 16, 11893–11900 (2025).

46 A. Mishra, J. Zubia-Aranburu, L. Zuzic, K. Saeed, J. Nielsen, B. Schiott, M. Dong, D. E. Otzen, and T. Weidner, “Structure of -synuclein bound to polystyrene surfaces probed by experimental and theoretical sum frequency generation spectroscopy,” Langmuir 42, 7653–7659 (2026).

47 L. J. Knight, F. N. F. Parker-Jurd, M. Al-Sid-Cheikh, and R. C. Thompson, “Tyre wear particles: an abundant yet widely unreported microplastic?” Environmental Science and Pollution Research 27, 18345–18354 (2020).

48 A. J. Nihart, M. A. Garcia, E. El Hayek, R. Liu, M. Olewine, J. D. Kingston, E. F. Castillo, R. R. Gullapalli, T. Howard, B. Bleske, J. Scott, J. Gonzalez-Estrella, J. M. Gross, M. Spilde, N. L. Adolphi, D. F. Gallego, H. S. Jarrell, G. Dvorscak, M. E. Zuluaga-Ruiz, A. B. West, and M. J. Campen, “Bioaccumulation of microplastics in decedent human brains,” Nature Medicine 31, 1114–1119 (2025).

49 F. Oosawa and S. Asakura, Thermodynamics of the Polymerization of Protein (Academic Press, 1975).

50 M. F. Bishop and F. A. Ferrone, “Kinetics of nucleation-controlled polymerization. a perturbation treatment for use with a secondary pathway,” Biophysical Journal 46, 631–644 (1984).

51 T. P. J. Knowles, C. A. Waudby, G. L. Devlin, S. I. A. Cohen, A. Aguzzi, M. Vendruscolo, E. M. Terentjev, M. E. Welland, and C. M. Dobson, “An analytical solution to the kinetics of breakable filament assembly,” Science 326, 1533–1537 (2009).

52 S. I. A. Cohen, M. Vendruscolo, M. E. Welland, C. M. Dobson, E. M. Terentjev, and T. P. J. Knowles, “Nucleated polymerization with secondary pathways. I. time evolution of the principal moments,” J. Chem. Phys. 135, 065105 (2011).

53 G. Meisl, J. B. Kirkegaard, P. Arosio, T. C. T. Michaels, M. Vendruscolo, C. M. Dobson, S. Linse, and T. P. J. Knowles, “Molecular mechanisms of protein aggregation from global fitting of kinetic models,” Nat. Protoc. 11, 252–272 (2016).

54 T. C. T. Michaels, A. Šarić, J. Habchi, S. Chia, G. Meisl, M. Vendruscolo, C. M. Dobson, and T. P. J. Knowles, “Chemical kinetics for bridging molecular mechanisms and macroscopic measurements of amyloid fibril formation,” Annual Review of Physical Chemistry (2018).

55 T. C. T. Michaels, D. Qian, A. Šarić, M. Vendruscolo, S. Linse, and T. P. J. Knowles, “Amyloid formation as a protein phase transition,” Nature Reviews Physics 5, 379–397 (2023).

56 A. K. Srivastava, J. M. Pittman, J. Zerweck, B. S. Venkata, P. C. Moore, J. R. Sachleben, and S. C. Meredith, “-amyloid aggregation and heterogeneous nucleation,” Protein Science 28, 1567–1581 (2019).

57 B. O. W. Elenbaas, L. Khemtemourian, J. A. Killian, and T. Sinnige, “Membrane-catalyzed aggregation of islet amyloid polypeptide is dominated by secondary nucleation,” Biochemistry 61, 1465–1472 (2022).

58 B. Frieg, M. Han, K. Giller, C. Dienemann, D. Riedel, S. Becker, L. B. Andreas, C. Griesinger, and G. F. Schröder, “Cryo-em structures of lipidic fibrils of amyloid-β (1-40),” Nature Communications 15 (2024), 10.1038/s41467-023-43822-x.

59 D. M. Vadukul, M. Papp, R. J. Thrush, J. Wang, Y. Jin, P. Arosio, and F. A. Aprile, “-synuclein aggregation is triggered by oligomeric amyloid-42 via heterogeneous primary nucleation,” Journal of the American Chemical Society 145, 18276–18285 (2023).

60 A. Röntgen, Z. Toprakcioglu, and M. Vendruscolo, “A42 promotes the aggregation of -synuclein splice isoforms <i>via</i> heterogeneous nucleation,” FEBS Letters 599, 2750–2767 (2025).

61 K. Araya, K. Nakamura, E. Ishibashi, T. Q. Noguchi, M. Kuragano, and K. Tokuraku, “Structural analysis of the asymmetric interaction between amyloid 42 and -synuclein: Amyloid 42 oligomers promote -synuclein aggregation while -synuclein inhibits amyloid 42 aggregation,” Biochemical and Biophysical Research Communications 809, 153506 (2026).

62 C. E. G. Leyns, A. Prigent, B. Beezhold, L. Yao, N. G. Hatcher, P. Tao, J. Kang, E. Suh, V. M. Van Deerlin, J. Q. Trojanowski, V. M. Y. Lee, M. E. Kennedy, M. J. Fell, and M. X. Henderson, “Glucocerebrosidase activity and lipid levels are related to protein pathologies in parkinson’s disease,” npj Parkinson’s Disease 9, 74 (2023).

63 G. Meisl, X. Yang, E. Hellstrand, B. Frohm, J. B. Kirkegaard, S. I. A. Cohen, C. M. Dobson, S. Linse, and T. P. J. Knowles, “Differences in nucleation behavior underlie the contrasting aggregation kinetics of the Aβ 40 and Aβ 42 peptides,” Proc. Natl. Acad. Sci. U.S.A. 111, 9384–9389 (2014).

