## Supplemental Information for "Lipidic *α*-synuclein fibrils catalyze the formation of pathological mature fibrils"

### SUPPLEMENTARY INFORMATION

#### EXPERIMENTAL METHODS

##### Protein production

N-terminally acetylated $\alpha$S was expressed in *E.coli* and purified as previously described (Hoyer2002. DOI: 10.1016/s0022-2836(02)00775-1). Briefly, BL21 pLysS cells were transformed with plasmids encoding *SNCA* and the pNatB acetylation complex. The expression was induced with IPTG, and the cells were lyzed with osmotic shock buffer. The expressed protein was purified, first by heat precipitation of endogenous protein, then ion exchange chromatography and finally size exclusion chromatography with phosphate buffer at pH 6.5 (20 mM $Na_2HPO_4$/$NaH_2PO_4$, 0.01% w/v sodium azide).

##### Formation of lipid model membranes

DMPS was purchased as a powder from Avanti Lipids. The lipid powder was resolubilized in a 9:1 chloroform:methanol mixture. The solvent was evaporated in a round-bottom flask using nitrogen gas to form a lipid film. The lipid film was re-suspended with the experimental buffer for 1.5 hours at 50°C. Finally, the suspension was extruded through polycarbonate membranes with a 100 nm pore diameter (Avanti Polar Lipids, Inc.). The size distribution of the resulting unilamellar lipid vesicles was confirmed by dynamic light scattering measurements.

##### Aggregation experiments

All aggregation experiments were measured under quiescent conditions in Corning 96-well Polystyrene Microplates (Corning 3601) on a BMG Labtech Clariostar microplate reader at 30°C. All samples contained 50 $\mu$M N-terminally acetylated $\alpha$S, unless stated otherwise, DMPS at varying concentrations as stated in the figures and legends, 50 $\mu$M ThT, and in 20 mM phosphate buffer pH 6.5.

##### Electron microscopy imaging

Once the samples had reached a plateau in aggregation, an aliquot was taken and applied to a carbon-coated grid. The grid was subsequently treated with 2% uranyl acetate for 30 s and then allowed to dry. Finally, excess stain was washed of the grid with $dH_2O$. Transmission electron microscopy images were acquired using a Philips CM100 transmission electron microscope at the Core Facility for Integrated Microscopy, Faculty of Health and Medical Sciences, University of Copenhagen.

#### SUPPLEMENTARY FIGURES

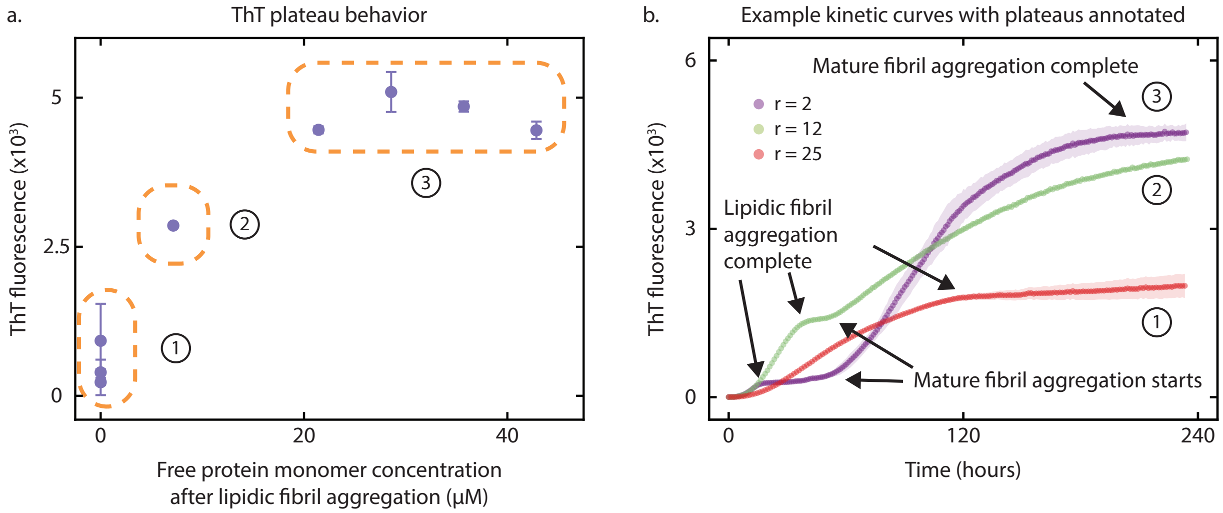


FIG. S1. **ThT plateau behavior when the initial concentration of lipids is varied.** **(a)** 1 - A consistent second growth phase is not observed in the timescale of the experiment. Increased noise in the observed plateau is likely due to the relatively low free monomer concentration available for nucleation of mature fibrils, leading to a stochastically determined lag phase for mature fibril aggregation. 2 - Plateau of $r = 12$ curve in the vary lipid dataset, which has not reached a steady-state ThT plateau within the timescale of the experiment (see green curve in panel (b)). 3 - Recorded ThT plateaus are constant, not scaling with free protein monomer remaining after the lipidic fibril aggregation reaction, for completed reactions as total protein monomer concentration is constant (providing further evidence for our proposed kinetic model containing a mechanism of converting lipidic fibrils to mature fibrils). **(b)** Example aggregation curves highlighting how plateau behavior has been grouped.

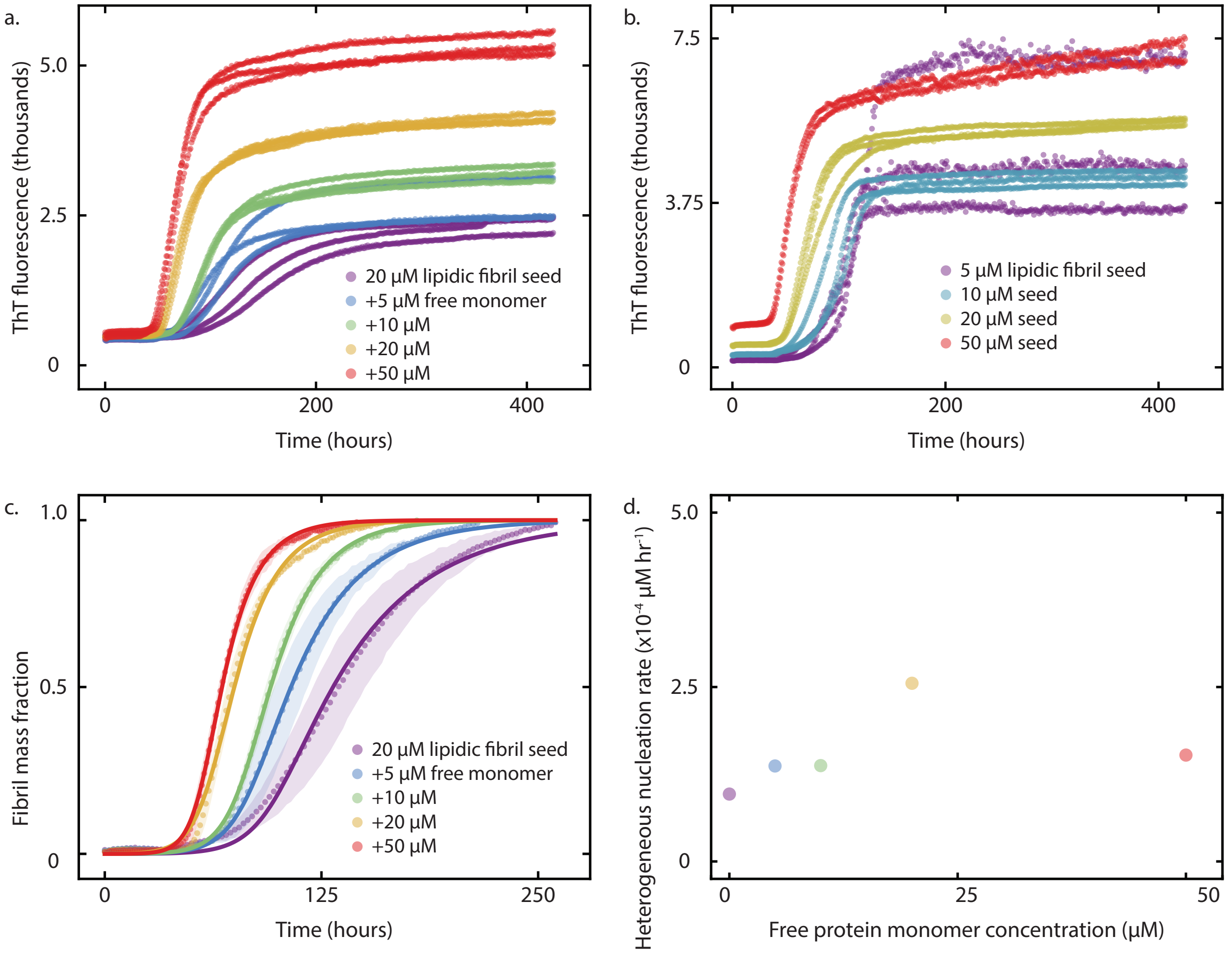


FIG. S2. **Effect of lipidic fibril seeds on mature $\alpha$-synuclein fibril aggregation.** **(a)** Raw ThT fluorescence kinetic profiles for lipidic fibril seeding experiment with increasing concentrations of free $\alpha$-synuclein monomer added at t = 0. **(b)** Raw ThT fluorescence kinetic profiles for lipidic fibril seeding experiment with increasing concentrations of lipidic fibril seed added at t = 0. **(c)** Individually fitting a coarse-grained primary nucleation rate in our theoretical model (solid lines), Eq. (6). **(d)** To capture the time evolution of mature fibril aggregation reveals that the primary nucleation rate is constant (within the same order of magnitude) as the free protein monomer concentration is increased.

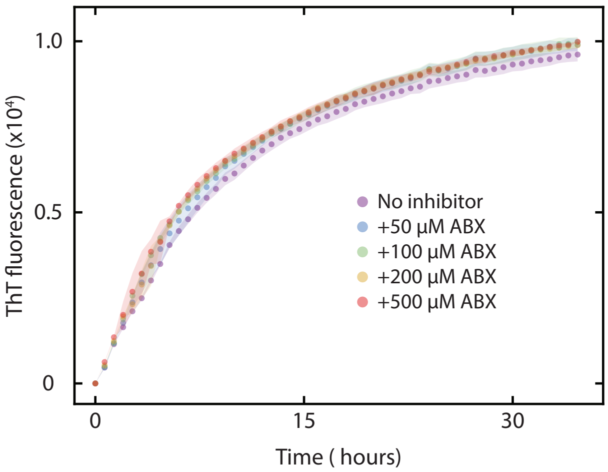


FIG. S3. **Presence of mature fibril seeds accelerates lipid-independent aggregation and is unaffected by the presence of ABX.** Mature fibril seeds, pre-formed in the absence of lipids in the PS plate, bypass the lag phase in lipid-independent $\alpha$-synuclein aggregation. Addition of the small-molecule ABX, known to inhibit lipid-induced aggregation to form lipidic fibrils, does not affect the kinetics of the seeded aggregation reactions.

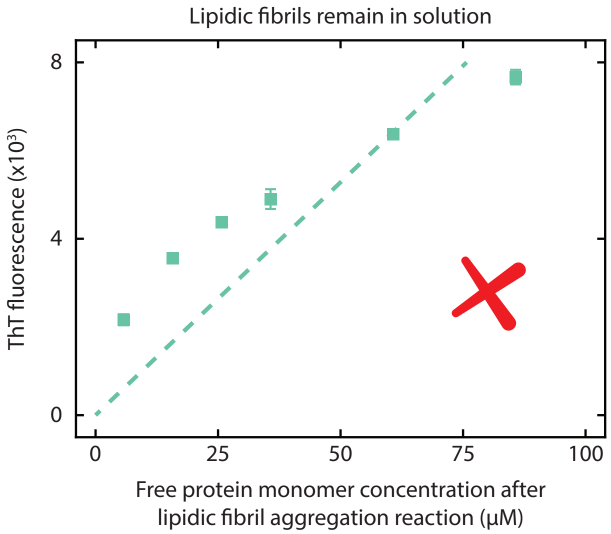


FIG. S4. **ΔThT does not directly scale with free protein monomer concentration remaining after lipidic fibril aggregation is complete.** Assuming that the protein monomer consumed to form lipidic fibrils is sequestered in this state and not available during the second growth phase does not accurately explain the scaling behavior observed, compared to assuming that all protein monomer is available for the second growth phase reaction (shown in Fig. 3c).

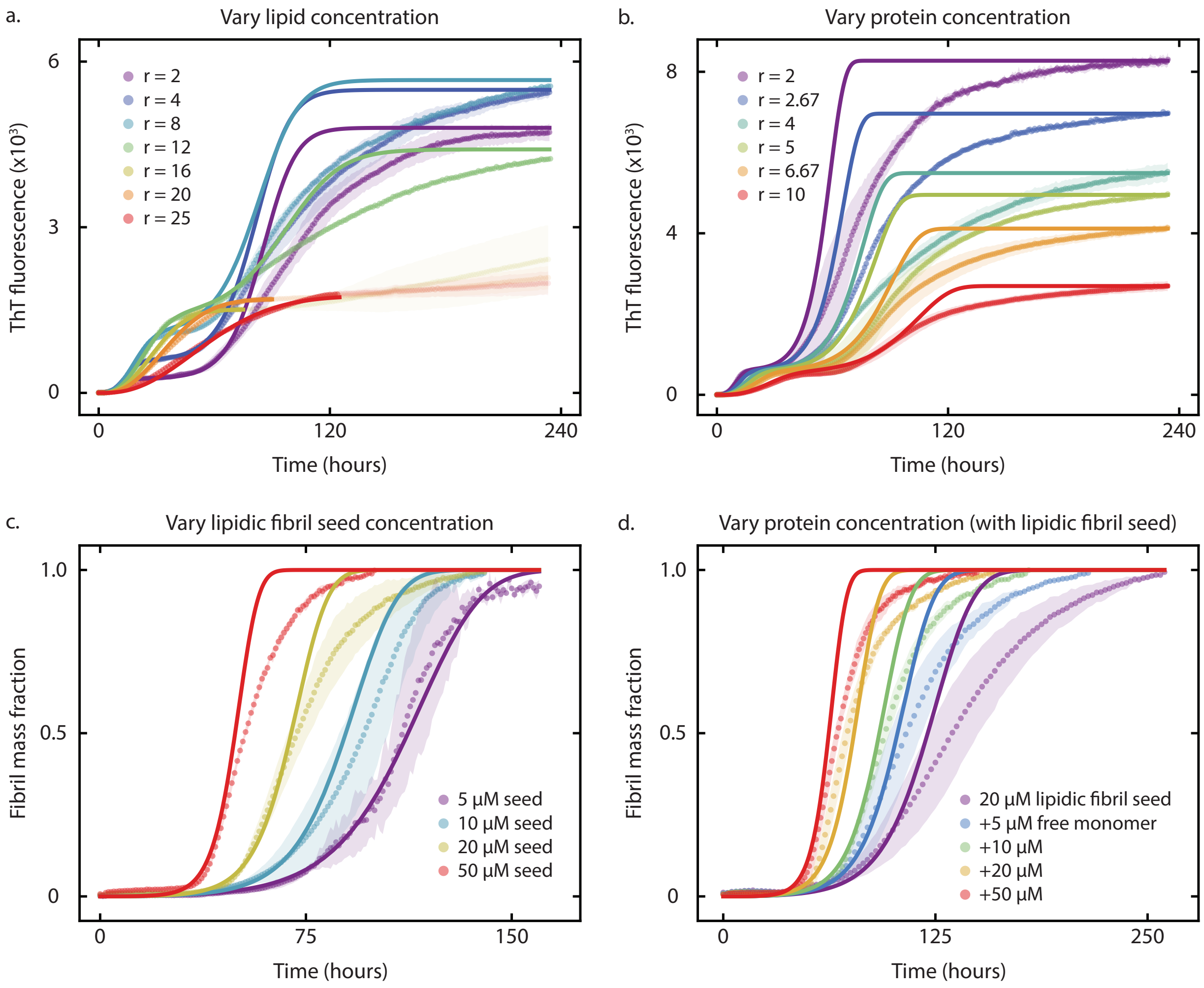


FIG. S5. **Misfits between theory and kinetic data highlight the role of lipidic fibril dissolution in controlling the timescale of mature fibril aggregation reaching completion.** Experimental ThT fluorescence measurements of the time evolution of *in vitro* lipid-dependent $\alpha$-synuclein aggregation, under varying initial concentrations of **(a)** lipid, **(b)** $\alpha$-synuclein monomer, **(c)** lipidic fibril seed and **(d)** $\alpha$-synuclein monomer in the presence of lipidic fibril seeds. All fitting parameters were kept consistent with those identified in the analysis of the experimental data in Fig. 3, 4 and Supplementary Material Fig. S2, only changing the $c$ parameter to match the definition used in kinetic models of homogeneous protein aggregation (see Methods).

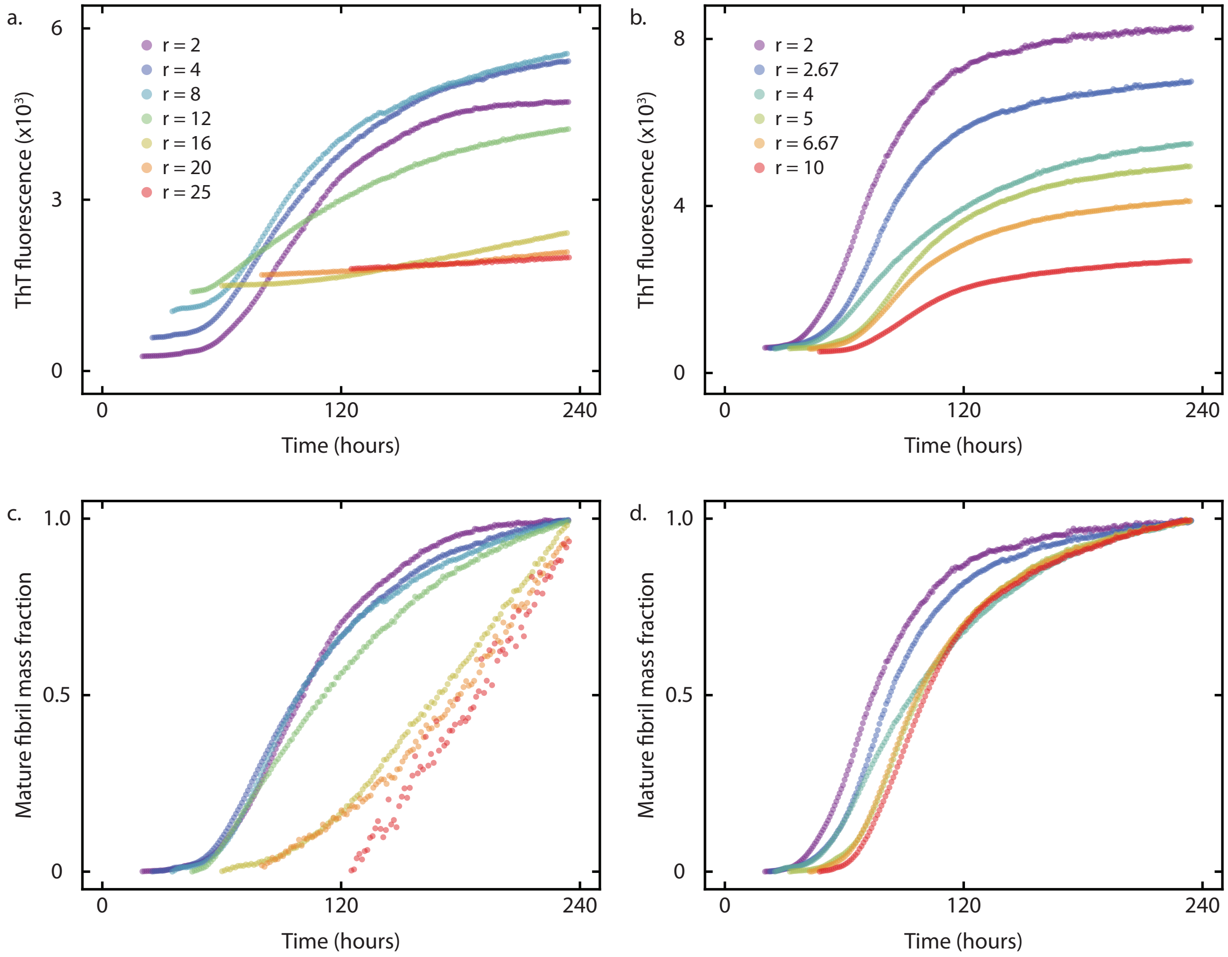


FIG. S6. **Global fitting procedure employed in this study to analyze mature fibril aggregation.** **(a)** Raw ThT fluorescence data capture the kinetics of mature fibril aggregation for varying initial lipid concentrations, where we segment the data to only include the intermediate plateau and second growth phase. **(b)** Raw ThT fluorescence data capture the kinetics of mature fibril aggregation for varying initial protein monomer concentrations. **(c)** The raw data in (a) are then individually normalized by the maximum fluorescence recorded. **(d)** The raw data in (b) are then individually normalized. Our theoretical model describing the kinetics of mature fibril aggregation is fitted to the normalized data, where $k_+k_h$, $k_+k_2$ and $c$ are globally fitted to the vary lipid dataset and then to the vary protein monomer dataset.

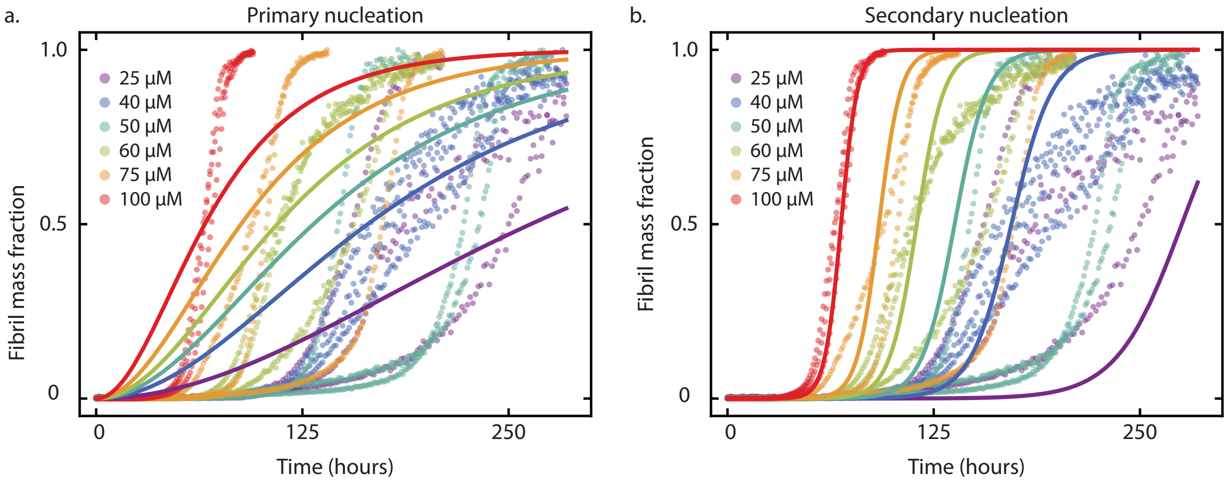


FIG. S7. **Lipid-independent $\alpha$-synuclein aggregation of mature fibrils in PS plate involves secondary nucleation.** Solid lines show model best fits of experimental ThT fluorescence data tracking the aggregation of varying initial $\alpha$-synuclein monomer concentrations to a kinetic model where fibrils can only form via primary nucleation **(a)** or primary and surface-catalyzed secondary nucleation **(b)**. Parameters for secondary nucleation fit: $\sqrt{k_+k_n} = 1.91 \times 10^{-6}\,\mathrm{hr}^{-1}\,\mu\mathrm{M}^{-n_c/2}$, $\sqrt{k_+k_2} = 1.43 \times 10^{-3}\,\mathrm{hr}^{-1}\,\mu\mathrm{M}^{-(n_2+1/2)}$, $n_c = 2$ and $n_2 = 1$. Details of the mathematical models are in Michaels *et al.*[55].

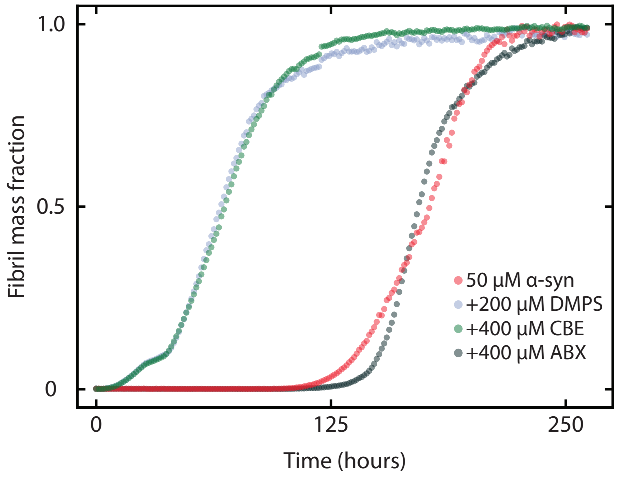


FIG. S8. **The small-molecule CBE has no inhibition effect on $\alpha$-synuclein aggregation.** At a concentration where ABX fully attenuates the aggregation of lipidic fibrils, and therefore the lipid-dependent aggregation pathway to form mature fibrils, CBE has no effect (comparing light blue to green curves).

#### SUPPLEMENTARY TABLES

TABLE I. Definitions of all symbols used in the theoretical modeling of the lipid-dependent aggregation of $\alpha$-synuclein.

| Symbol | Definition |
|---|---|
| $m_{\mathrm{tot}}$ | Total initial concentration of protein monomers |
| $L_{\mathrm{tot}}$ | Total initial concentration of lipids |
| $M_{lf}$ | Mass concentration of lipidic fibrils |
| $M_{mf}$ | Mass concentration of mature fibrils |
| $\beta$ | Average number of lipid units bound by a single protein monomer |
| $\alpha$ | Average number of lipid units bound by a single protein monomer in fibril form |
| $k_{\mathrm{on}}$ | Rate of free monomer binding to the lipid surface |
| $k_{\mathrm{off}}$ | Rate of bound monomer dissociation from the lipid surface |
| $K_{\mathrm{D}}$ | Dissociation constant of protein-lipid binding |
| $k'_{+}$ | Rate constant describing lipidic fibril elongation |
| $k_o$ | Rate constant describing primary nucleation of protein oligomers |
| $k_c$ | Rate constant describing the conversion of protein oligomers to lipidic fibrils |
| $n_1$ | Reaction order parameter describing the dependence of primary nucleation on bound monomers |
| $n_2$ | Reaction order parameter describing the dependence of primary nucleation on free monomers |
| $n_l$ | Reaction order parameter describing the dependence of the reaction flux of oligomer conversion on total lipid concentration |
| $\theta_0$ | Approximated solution for the surface coverage of protein monomers applicable at early-time scales |
| $t_{1/2}$ | Time for half the total concentration of protein monomers to convert to fibril form |
| $\tau_c$ | Timescale for conversion of oligomers to fibrils |
| $k_{+}$ | Rate constant describing mature fibril elongation |
| $k_n$ | Rate constant describing polystyrene-induced primary nucleation of mature fibrils |
| $k_h$ | Rate constant describing heterogeneous primary nucleation of mature fibrils |
| $k_2$ | Rate constant describing secondary nucleation of mature fibrils |
| $n_c$ | Reaction order parameter describing the dependence of polystyrene-induced primary nucleation on protein monomers |
| $n_h$ | Reaction order parameter describing the dependence of heterogeneous primary nucleation on protein monomers |
| $n_s$ | Reaction order parameter describing the dependence of secondary nucleation on protein monomers |
| $K_{M,h}$ | Saturation constant for heterogeneous primary nucleation of mature fibrils |
| $K_{M,s}$ | Saturation constant for secondary nucleation of mature fibrils |

TABLE II. Summary of fitting parameters used to describe the experimental data in Fig. 4

| Parameter | Vary lipid concentration (Fig. 4a) | Vary protein concentration (Fig. 4b) |
|---|---|---|
| $\sqrt{k'_+ k_o}$ | $0.101\,\mathrm{hr}^{-1}\,\mu\mathrm{M}^{(1-n_2)/2}$ | $0.239\,\mathrm{hr}^{-1}\,\mu\mathrm{M}^{(1-n_2)/2}$ |
| $k_c$ | $1.22\times10^{-2}\,\mathrm{hr}^{-1}\,\mu\mathrm{M}^{-n_l}$ | $0.509\times10^{-2}\,\mathrm{hr}^{-1}\,\mu\mathrm{M}^{-n_l}$ |
| $\alpha$ | 16 | 16 |
| $n_1$ | 0 | 0 |
| $n_2$ | 1.5 | 1.5 |
| $n_l$ | 0.321 | 0.103 |
| $\sqrt{k_+ k_h}$ | $4.91\times10^{-4}\,\mathrm{hr}^{-1}\,\mu\mathrm{M}^{-(1+n_h)/2}$ | $6.97\times10^{-4}\,\mathrm{hr}^{-1}\,\mu\mathrm{M}^{-(1+n_h)/2}$ |
| $\sqrt{k_+ k_2}$ | $1.06\times10^{-2}\,\mathrm{hr}^{-1}\,\mu\mathrm{M}^{-1/2}$ (multiplied by $m_{\mathrm{tot}}^{n_s}$) | $0.944\times10^{-2}\,\mathrm{hr}^{-1}\,\mu\mathrm{M}^{-1/2}$ |
| $n_h$ | 0 | 0 |
| $n_s$ | 0.586 | 0 |
| $c$ | 0.257 ($L_{\mathrm{tot}} = 100\,\mu\mathrm{M}$)<br>0.222 ($L_{\mathrm{tot}} = 200\,\mu\mathrm{M}$)<br>0.242 ($L_{\mathrm{tot}} = 400\,\mu\mathrm{M}$)<br>0.300 ($L_{\mathrm{tot}} = 600\,\mu\mathrm{M}$) | 0.195 ($m_{\mathrm{tot}} = 100\,\mu\mathrm{M}$)<br>0.199 ($m_{\mathrm{tot}} = 75\,\mu\mathrm{M}$)<br>0.207 ($m_{\mathrm{tot}} = 50\,\mu\mathrm{M}$)<br>0.258 ($m_{\mathrm{tot}} = 40\,\mu\mathrm{M}$)<br>0.368 ($m_{\mathrm{tot}} = 30\,\mu\mathrm{M}$)<br>0.645 ($m_{\mathrm{tot}} = 20\,\mu\mathrm{M}$) |
